# An ancient gene duplication is implicated in virulence in the human pathogen, *Histoplasma*

**DOI:** 10.64898/2026.02.25.708051

**Authors:** Victoria E. Sepúlveda, Jingbaoyi Li, David A. Turissini, Jonathan A. Rader, Oliver Kompathoum, Daniel R. Matute

**Affiliations:** Department of Biology, University of North Carolina at Chapel Hill, 27599 Chapel Hill, NC, USA; Department of Biology and Chemistry, Texas A&M International University, 78041 Laredo, Laredo, TX, USA

## Abstract

*Histoplasma* spp. is a dimorphic fungal primary pathogen that infects people worldwide and frequently affects immunosuppressed patients. Previous studies have identified the *AMY1* gene product, the α-amylase Amy1p, as essential for α-glucan production and virulence in *Histoplasma capsulatum*. We identified two new genes (*AMY2* and *AMY3*) in the *Histoplasma* genome that encode putative α-amylases and made mutants using CRISPR/Cas9 technology, followed by evaluation of their role in α-glucan biosynthesis and virulence. We also searched for *AMY* gene copies in 19 fungal genomes with the goals of identifying orthologs for *AMY2* and *AMY3*, and establishing how many *AMY* copies existed across different fungi. We found that the number and type of α-amylases vary depending on the fungal species; that all α-amylases related to *Histoplasma* Amy1p belong to the GH13_5 subfamily, and all orthologs related to *Histoplasma*’s Amy2p and Amy3p belong to the GH13_1 subfamily. We performed phylogenetic analyses of the three paralogs and revealed that the *Histoplasma AMY* duplications are ancient. We further established Amy2 is an ortholog of *Aspergillus niger* AgtA, and *Aspergillus nidulans* AmyD, and that it is partially involved in *Histoplasma* α-glucan biosynthesis and virulence, while Amy3p is an ortholog of *Aspergillus flavus* Amy1, and it is dispensable for α-glucan biosynthesis and virulence.

## INTRODUCTION

α-Amylases are a family of enzymes that catalyze the hydrolysis of starch and glycogen into simple sugars through the cleavage of α-(1,4)-glycosidic linkages. These enzymes are broadly distributed among microorganisms, plants, and animals, and play an essential role in carbohydrate digestion and metabolism. They constitute the Glycoside Hydrolase (GH) Family 13; this family of enzymes is further divided into 35 subfamilies based on substrate specificity and protein sequence (Stam et al. 2006). Fungal α-amylases were thought to be restricted to extracellular enzymes classified into the GH13_1 subfamily until a phylogenetic and biochemical study identified a novel cluster of intracellular fungal α-amylases, GH13_5, which, based on sequence homology, included the *Histoplasma* Amy1p (Van Der Kaaij, Janeček, et al. 2007). More recently, an *in silico* analysis of fungal α-amylases described members from subfamilies GH13_5 and GH13_32 to be evolutionarily more closely related to each other than to their counterparts from GH13_1 (Janíčková and Janeček 2020).

α-Amylases are implicated in fungal virulence in a variety of plant and animal pathogens (reviewed in (Rafiei et al. 2021)). One of these fungi is the dimorphic fungus *Histoplasma*, the causal agent of Histoplasmosis, one of the most common fungal respiratory diseases worldwide, with hundreds of thousands of new infections occurring annually (Ajello 1968; Wheat 2003; Kauffman 2007; Sepúlveda et al. 2024). Genetic mapping and functional assays have revealed genes required for virulence in *Histoplasma* (reviewed in (Sepúlveda et al. 2024)), *AMY1* (Amy1p) being one of them. The Amy1p enzyme is highly conserved in fungi and animals, and genetic comparisons revealed that the catalytic residues involved in the hydrolysis of α-1,4 glycosidic linkages are conserved in the *AMY1* orthologs of all ascomycetes, a group that diverged over 150 million years ago (Shen et al. 2020). In *Histoplasma*, Amy1p is an α-amylase of 540 amino acids involved in starch degradation. The *Histoplasma AMY1* gene product has all the predicted catalytic residues as other α-(1,4)-amylases, and is involved in synthesizing the primers α-1,4-linked glucooligosaccharide, which are required to create longer polymers and stabilize α-1,3-glucan *in vivo* (Grün et al. 2005; Yoshimi et al. 2017). At environmental temperatures of ∼25°C, *Histoplasma* grows as multicellular mold, forming hyphae and spores which typically do not express *AMY1*. At 37°C, *Histoplasma* transforms to its yeast form and expresses *AMY1,* which is required for the production of α-(1,3)-glucan, an adhesion molecule that allows evasion of host immune activity (Rappleye et al. 2007). For this reason, Amy1p is considered a virulence factor in *Histoplasma* (Marion et al. 2006). Previous efforts have shown that deletion of *AMY1* abolishes cell wall α-(1,3)-glucan and attenuates virulence (Marion et al. 2006). The loss of *AMY1* dramatically reduces the ability of *Histoplasma capsulatum sensu stricto* to kill macrophages and to colonize murine lungs.

*AMY1* also influences virulence in other fungi. In *Aspergillus nidulans*, the deletion of AmyG, an ortholog of Amy1, leads to reduced production of α-(1,3)-glucan (Camacho et al. 2012; Kazim et al. 2021). Introducing an *AMY1* copy from *Paracoccidioides brasiliensis* restores α-(1,3)-glucan production in an *AMY1*-null strain of *H. capsulatum,* and generates short oligosaccharides that act as primers at the first step of α-(1,3)-glucan biosynthesis (Camacho et al. 2012). *AMY1* shows higher expression in spherules than in mycelia in *Coccidioides immitis* (Viriyakosol et al. 2013).

Despite the conclusive evidence that Amy1 is involved in virulence in *Histoplasma* and other pathogenic fungi, there has been no formal test of the involvement of other Amys in virulence in *Histoplasma* or on any other fungi. Here we report that *Histoplasma* has two copies of α-amylases (Amy2p, Amy3p) that belong to the GH13_1 subfamily in addition to the α-amylase Amy1p, which belongs to the subfamily GH13_5. We detected no copies of GH13_32 which have been mostly detected in basidiomycetes. These results indicate that the Amy repertoire in *Histoplasma* is similar, but not always identical, to that of other ascomycetes, including several human pathogens. We thus evaluated the contribution of each of these amylases to α-(1,3)-glucan synthesis by evaluating colony morphology and virulence. All genes contribute to gene morphology but they do not always do so in the same direction.

While abrogating *AGS1*, *AMY1*, and *AMY2* leads to smoother colonies, disruption of *AMY3* has the opposite effect. Experimental macrophage infections confirmed that *AGS1* and *AMY1* are necessary for virulence. *AMY2* is partially required for macrophage infections but *AMY3* has no detectable contribution to virulence. Our results indicate that the dissection of the contributions of genes within gene families might be illuminating to understand how virulence evolves and is maintained in pathogens.

## MATERIALS AND METHODS

### Identification of AMY duplications and in silico analysis

#### Identification of *AMY* duplications

Fungal α-amylases belong to the glycoside hydrolase GH13 family, and some fungal genomes contain paralogs that arose through gene duplication. Because the only α-amylase described to date in *Histoplasma capsulatum* is Amy1p, which is critical to produce α-(1,3)-glucan, we searched for evidence of *AMY* gene duplication in *Histoplasma*. To do so, we looked for genes that had been annotated in the genome of *H. capsulatum* encoding putative glycoside hydrolases in addition to *AGS1* and *AMY1* by searching on FungiDB (https://fungidb.org/fungidb/app) for “alpha amylase”, filtering by organism, and using the *Histoplasma capsulatum* G186AR genome. We found only three candidate genes. The first one was *AMY1* (17150_01352), and two additional genes *AMY2* (17150_02300), and *AMY3* (17150_07038).

These three *H. capsulatum* genes were then protein blasted with default parameters (Blast V2.17.0, (Altschul et al. 1990)) against the G186AR *H. capsulatum* reference genome (Voorhies et al. 2022) to identify *AMY1*, *AMY2*, and *AMY3* from the same reference genome. The three G186AR *AMY* genes were then protein blasted with default parameters (Blast V2.17.0) against protein sequences from 19 genomes (Table S1). The protein sequences from these genomes were prepared for all annotated transcripts using python scripts that processed gff3 files downloaded from NCBI (Table S1). The blast hits with E-values < 1 × 10^-50^ were considered significant, and these genes were assigned to *AMY1*, *AMY2*, or *AMY3* based on which had the lowest E-value. Thirty-six known GH13 genes (Table S2) were also protein blasted against the protein sequences from the 19 genomes using Blast V2.17.0 with default parameters. To link the *AMY1*, *AMY2*, and *AMY3* names with species-specific naming conventions, the three G186A *H. capsulatum AMY* genes were blasted (Blast V2.17.0) using default parameters against eight *AMY* genes from *Aspergillus flavus*, *Aspergillus fumigatus*, *A. nidulans*, *Aspergillus niger*, and *P. brasiliensis*.

#### Protein alignment and BLAST

We used the full-length Amy1p, Amy2p, and Amy3p sequences from *H. capsulatum sensu stricto* G186A, and the concatenated sequences corresponding to the four conserved regions comprising the catalytic sites of the GH13 family of α-amylases, to determine the percentage of similarity among the three Amy proteins by BLAST analysis using the NCBI protein BLAST tool. We performed protein alignments with the bioinformatics software Geneious Prime using the Clustal Omega default parameters.

#### *AMY* chromosomal arrangement and population nucleotide diversity

We used nucleotide Blast v2.17.0 to identify the locations of *AMY1*, *AMY2*, and *AMY3* within five highly contiguous *Histoplasma* genomes (Voorhies et al. 2022). Blast results with the lowest E-values were selected. We used the positions and relevant chromosomes of the three *AMY* genes to generate a chromosomal plot using the R library Chromomap v4.1.1 (Anand and Rodriguez Lopez 2022). We also calculated nucleotide diversity, π, using Pixy v1.2.11 (Korunes and Samuk 2021) to compare nucleotide diversity across species of *Histoplasma*. We used a previously generated Variant Call File (VCF) from (Ly et al. 2025; Mapengo et al. 2025) as input to Pixy, where π was calculated in 100bp windows. Genomic averages of π were calculated per species using the mean() function in R. The locations of *AMY1*, *AMY2*, and *AMY3* were identified in the reference genome of *H. capsulatum* isolate H88 (accession GCA_017310615.1) in correspondence with the reference used within the VCF.

#### Detection of signal peptides and transmembrane domains

Using full-length protein sequences, we searched for three types of sequences that determine cellular location. First we looked for secretion signal peptides using SignalP 6.0 (Teufel et al. 2022). Second, we screened for putative glycosylphosphatidylinositol (GPI)-anchor signal sequences using the bioinformatic prediction tool NEtGPI v1.1 (Gíslason et al. 2021). Finally we assessed for the presence of transmembrane domains using TMHMM 2.0 (Krogh et al. 2001).

### CRISPR mutants

#### Fungal strains and culture conditions

We used the wild-type *H. capsulatum sensu stricto* Panamanian isolate G186A (ATCC 26029, same as G186AR) to generate all mutants used in this study. *Histoplasma capsulatum* yeasts were grown in *Histoplasma* Macrophage Medium (HMM, solid or liquid) as previously described (Worsham and Goldman 1988) at 37°C with 95% air-5% CO_2_. Liquid cultures were grown with continuous shaking (150 rpm in a Multitron orbital shaker, Infors HT). Solid medium contained 0.6% agarose (SeaKem ME grade) and 25 mM FeSO_4_.

#### Plasmid design and transformation of *E. coli*

We generated plasmids to disrupt the genes encoding *AGS1, AMY1, AMY2*, and *AMY3* using CRISPR technology optimized for *Histoplasma* (Rappleye 2023) to assess the role of each new putative α-amylase in the production of α-(1,3)-glucan. Single guide RNA (sgRNA) consists of the CRISPR RNA (crRNA) and trans-activating crRNA (tracrRNA) in one molecule, while dual/tandem guide RNAs (tgRNA) use two distinct gRNA cassettes simultaneously. Every gRNA cassette consists of 6 nucleotides reverse-complementary to the first 6 nucleotides on the 5’ end of crRNA, which allows RNA duplex formation and ribozyme-based cleavage, followed by a hammerhead ribozyme sequence, a 20-nucleotide crRNA sequence, and a gRNA scaffold (as described in (Rappleye 2023). We generated 3 mutants for each gene using two different sgRNAs and a tgRNA (Table S3). First, we used a sgRNA targeting the 5’ coding sequence (CDS) region of each gene. Second, we used a sgRNA targeting the 3’ CDS region of the gene. All sgRNA introduced a frameshift indel mutation (Table S3). Finally, we used a tgRNA designed to delete a portion of the gene located between the two sgRNAs (at least 1Kb for each gene, Table S3).

We used CRISPOR ((Concordet and Haeussler 2018), https://crispor.gi.ucsc.edu/crispor.py) to design gRNAs against *AGS1, AMY1, AMY2,* and *AMY3*, using the coding gene sequence as input and selecting “*Histoplasma* strain G217B” as the selected genome and “20bp-NGG-SpCas9, SpCas9-HF1, eSpCas9 1.1” as the Protospacer Adjacent Motif from the dropdown menus. We selected candidate protospacers with high prediction efficiency scores: Wang scores (>70), Doench “16 scores” (>65), and Chari scores (>60) (Table S3). We used Twist Biosciences (San Francisco, California) to synthesize CRISPR RNAs (crRNAs, which include the selected candidate protospacers targeting genes of interest (crRNA components are described in detail in (Rappleye 2023)). For sgRNAs, we used the In-Fusion Snap Assembly Master Mix kit (Takara Bio) to clone the crRNAs into the SwaI-digested CRISPR/Cas9 telomeric vector (kindly provided by Dr. Chad Rappleye). We transformed 10 ng CRISPR plasmids into chemically competent *Escherichia coli* HST08 Stellar Competent Cells (Takara Bio) by heat shocking for 45 seconds at 42 ℃. Transformed cells were allowed to recover in non-selective liquid LB media for 45 minutes before plating on selective LB agar supplemented with kanamycin (50 ug/mL). For tgRNA vectors, we digested the two sgRNA vectors targeting the same gene with restriction enzymes: one with *Hind*III and *Swa*I and the other with HindIII and *Stu*I, and gel-purified the fragments containing the gRNAs. Fragments were cloned together using T4 DNA ligase (New England Biolabs), followed by electroporation into *E. coli* DH5ɑ (2.5kV, 600Ω, 25 uF). Transformed cells were allowed to recover in non-selective liquid LB media for 45 minutes before plating on selective LB agar supplemented with kanamycin (50 ug/mL).

#### Electroporation of *Histoplasma*

We electroporated *Histoplasma* cells using protocols previously described (Woods et al. 1998). We grew *Histoplasma* yeast cells to exponential phase, performed a low-speed centrifugation for 3 minutes at 600 rpm (Allegra X-30, Beckman centrifuge) to remove large yeast clumps, and transferred the supernatant to a new tube to collect single cells by centrifugation for 5 minutes at 1800 rpm. We washed the pellet with 10% mannitol and resuspended it in 200 ul of 10% mannitol. We electroporated *Histoplasma* yeast cells (0.75kV, 600Ω, 25 uF) using 100-200 ng of linearized plasmid (*Pac*I-digested), then transferred cells to 5ml of non-selective media (HMM) and allowed them to recover for 6-12 hours before plating them on selective media containing hygromycin B (200 μg/ml). Putative mutants were screened by PCR. We designed primers flanking the potential deletion in each target gene. Using the DNA of putative mutants as templates, we amplified the genomic region for each gene and ran DNA gels to visualize PCR products. PCR products of tgRNA mutants are predictable and are smaller than the WT’s PCR products due to deletion of at least 1kb of the WT sequence (Table S3). For sgRNA mutants, PCR products were Sanger sequenced to confirm mutations.

### Phenotypic assays

We studied the phenotypic characteristics of each produced mutant. Since *H. capsulatum AGS1* and *AMY1* have been previously implicated in colony morphology (Marion et al. 2006; Rappleye et al. 2007), where wild type (WT) and α-glucan producing-strains are characterized by a rough colony morphology, and smooth colonies are a proxy for lack of α-glucan production we evaluated colony morphology.

#### Colony morphology

We grew *H. capsulatum cultures* (WT and mutants) to late exponential phase and harvested and washed them with HMM broth. We prepared serial dilutions (1×10^7^, 1×10^6^, 1×10^5^, 1×10^4^, 1×10^3^, and 1×10^2^) and spotted drops of 10 uL of each yeast serial dilution onto HMM plates. We incubated for 7 days at 37°C and took pictures to assess colony morphology. Colonies were categorized as rough or smooth. To ensure reproducibility, we examined at least six colonies per strain and confirmed no isolate showed variation in colony morphology. We further quantified colony rugosity for each genotype using a custom image analysis pipeline implemented in the image analysis toolbox in Matlab (MATLAB 2025). First, we generated a binary mask of the raw image using an edge detection method that was programmed to detect circular (or nearly circular) objects within a diameter range bracketing the pixel diameter of the colonies in the images. Each raw image was then converted from RGB colorspace to 8 bit grayscale. We further applied an entropy filter to the grayscale image with the function ‘entropyfilt’ (Gonzalez 2009). This filter calculated a value of image complexity (or pixel randomness) for a 9×9 pixel neighborhood around each pixel in the image. Briefly, entropy values vary directly with colony rugosity; smooth colonies have relatively low entropy and rough colonies have higher entropy values. We applied the binary mask to the entropy-filtered image, segregating each of the colonies within the image. Pixel areas for each colony were approximately 40,000. We randomly sampled 1,000 pixels from each colony and extracted the calculated entropy values for each. The mean pixel entropy values of the WT varied slightly across the four analyzed images, so we used a z-score standardization to rescale the entropy values of each image such that the WT mean was 0 and standard deviation was approximately 1, facilitating direct comparison across images. We used Wilcoxon rank sum tests (R function *wilcox.test* (library *stats*, (R Core Team 2018)) to compare the mean entropy values between colonies of wild-type and mutant strains.

#### Growth Curves

We measured the growth rate of five genotypes of *H. capsulatum* (WT and four mutants generated using tgRNA) in liquid media. For the growth curves, we inoculated 35 mL of HMM broth at a final concentration of 1 × 10^6^ yeast/mL and grew the culture for 9 days. We removed 600 μL from each culture (or media to use as blanks) and mixed them with 300 μL of 3M NaOH in a plastic cuvette, which was covered with Parafilm (Pechiney plastic packaging) and vortexed for 10 seconds to separate yeast clumps. We measured optical density (OD) using a GENESYS™ 10 UV-Vis Spectrophotometer (Thermo Scientific) starting at Day 0 and again every 24 hours thereafter until Day 9. To study the parameters of the growth curve, we followed the same approaches described in Sepúlveda et al. (2024, May 21). Growth curves were performed at least twice per genotype. To quantify the rate of growth, we used a four-parameter logistic model with the formula:

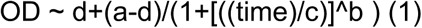

where *a* is the OD at the beginning of the experiment (presumably close to zero), *b* is the rate of increase in OD at point *c*, the inflection point of the curve, and *d* is the maximum OD in the curve, the asymptote. This model allows for an initial growth phase in which cells divide without increasing OD (lag-phase) and incorporates an asymptote, estimated from the data, at which cells cease replication or at which OD stops showing significant change. We tested 10 starting values per parameter and selected the model with the lowest Akaike Information Criterion (AIC; (Akaike 1973)) using the function AIC (library *stats*; (R Core Team 2018)). We fit a non-linear regression for each genotype. To determine whether the four fitted parameters differed among genotypes, we generated 1,000 bootstrap regressions using the R function nls.boot (library *nlstools*; (Baty et al. 2013; Baty et al. 2015)). We then compared the values of *a*, *b*, *c*, and *d* among the five genotypes using non-parametric tests (Wilcoxon rank-sum test with continuity correction; function *wilcox.test*, library *stats*; (R Core Team 2018)).

#### Macrophage virulence assays

Finally, we studied the ability of each genotype to penetrate and kill host cells. We cultured P388D1 macrophage-like cells in F-12 medium (Life Technologies) with 10% fetal bovine serum (HyClone) and 1% L-glutamine or in HMM-M when coincubated with *Histoplasma* yeast cells (Eissenberg et al. 1988; Eissenberg et al. 1991). We followed protocols previously described to determine *Histoplasma* virulence in P388D1 cells (Sebghati et al. 2000). We infected monolayers of 1 × 10^5^ P388D1 cells with WT and mutant yeasts (at least 3 replicates per genotype) at a multiplicity of infection of 1:3 (yeasts: macrophages) in 24- well cell culture plates (Corning Life Sciences) and incubated them at 37°C in 95% air–5% CO_2_ on a nutator.

We replaced fifty percent of the culture medium with fresh medium every other day, starting on Day 3. After 6 days of infection, we removed the liquid culture medium and yeast cells, and lysed the remaining macrophages with macrophage lysis buffer (20 mM Tris, 5 mM EDTA, 0.05% SDS). We used the PicoGreen double-stranded DNA quantification reagent (Molecular Probes) to measure the amount of macrophage DNA in each well by fluorescence spectroscopy (Excitation = 480nm, emission = 520nm) on a 96-well round-bottom plate (Fisher Scientific) using a Synergy HT plate reader (Biotek Instruments, Winooski, VT). We modeled the data using the function *lm* (library stats, (R Core Team 2018)): each genotype was the fixed effect and the proportion of remaining viable macrophages was the response. We used the R function *anova* (library stats, (R Core Team 2018)) to determine significance of the genotype effect. To compare the means between genotypes, we used a Tukey post-hoc test as implemented in the R function *glht* (library *multcomp*, (Hothorn et al. 2012; Hothorn et al. 2016)). We also evaluated whether the sgRNA mutations and the tgRNA mutations showed similar levels of virulence in macrophage experiments. To do so, we fit a factorial linear model in which the number of remaining macrophages was the response, and the gene (*AGS1* and *AMY1*) and the type of mutation were fixed effects. The model also included the interaction between these two effects. We used the R function *lm* (library *stats*, (R Core Team 2018)) to fit the model and *anova* (library *stats*, (R Core Team 2018)) to assess the significance of the fixed effects.

## RESULTS

### Identification and *in silico* analysis of an α-amylase gene family in *Histoplasma*

In addition to *AMY1* and *AGS1*, we found evidence of two genes encoding putative glycoside hydrolases in the genome of *H. capsulatum* strain G186AR: HCBG_00468 and HCBG_03431. These genes are unlinked in the genome (Figure 1A) and differ in their sequence (Figure 1B). We named these genes *AMY2* (HCBG_00468) and *AMY3* (HCBG_03431). While Amy1p has 540 amino acids, the predicted Amy2p and Amy3p have 473 and 500 aa, respectively. The full length-protein sequences indicate that Amy2p has 21.73 % identity to Amy1p, while Amy3p has 28.42 % identity. The sequence comparison between Amy2p and Amy3p shows a 41.18 % identity. The GH13 family of α-amylases is characterized by four conserved regions comprising the catalytic sites (MacGregor et al. 2001). Although the similarity between the full-length protein sequences of these three α-amylases is below 50 %, alignment and comparison of these conserved regions showed the identity between Amy1p and Amy2p as well as Amy1p and Amy3p increased to 62.96 %, while the % identity increased to 76.92 % in the case between Amy2p and Amy3p (Figure 1B). Alignment of the Amy1p sequence with known α-(1,4)-amylases from fungal pathogens *A. fumigatus, Cryptococcus neoformans, Blastomyces dermatitidis, P. brasiliensis, Emmonsia crescens,* and *C. immitis*, and the non-pathogenic *Neurospora crassa* shows all essential catalytic residues are conserved in *H. capsulatum* (Figure 2A). These acidic residues essential for catalysis, a hallmark of the GH13 family of α-amylase, and present in *H. capsulatum* Amy1p (D266, E296, and D362, Figure 2A) are also present and conserved in the two Amy1p paralogs Amy2p and Amy3p (Figure 2B). Additionally, the lysine and histidine residues within the conserved region II (Amy1p 262_GLRFDAA**KH**), linked with hydrolysis of α-(1-4)-glycosidic bonds, were also present in Amy2p and Amy3p (Figure 2B). These analyses suggest that the two newly identified Amy1p paralogs in *Histoplasma* are members of the GH13 family of α-amylase. Notably, the three genes show exceptionally low levels of polymorphism (Figure 2E-G) compared to the rest of the genome.

**FIGURE 1.**
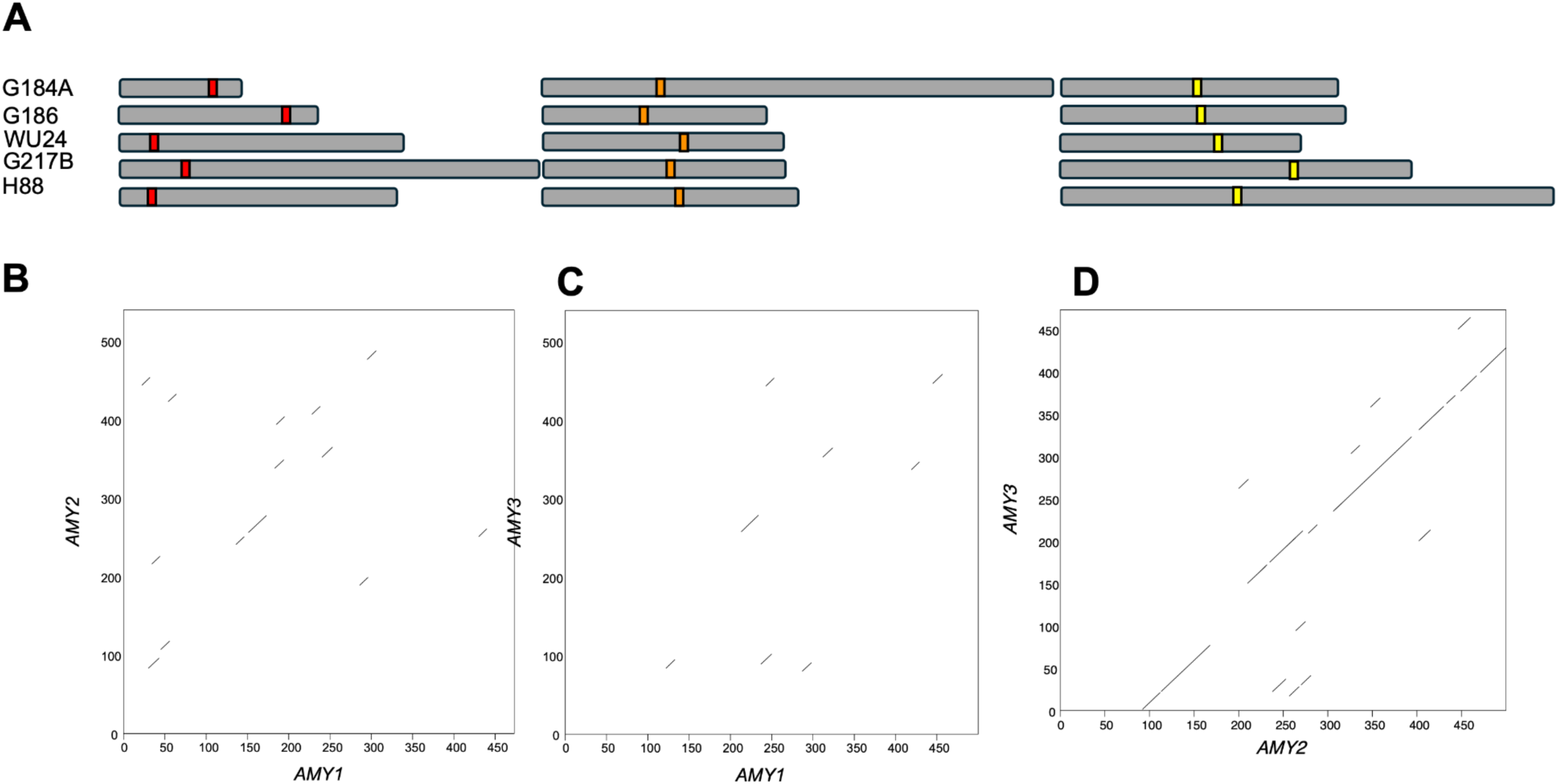
The *Histoplasma capsulatum ss* genome harbors three putative α-amylases. **A.** The three genes are unlinked in the genome in four species of *Histoplasma*. **B-D.** Dotplots comparing the three copies of the three putative α-amylases. Red: *AMY1*, orange: *AMY2*, yellow: *AMY3*. **B.** Amy1p vs Amy2p. **C.** Amy1p vs Amy3p. **D.** Amy2p vs. Amy3p.

**FIGURE 2.**
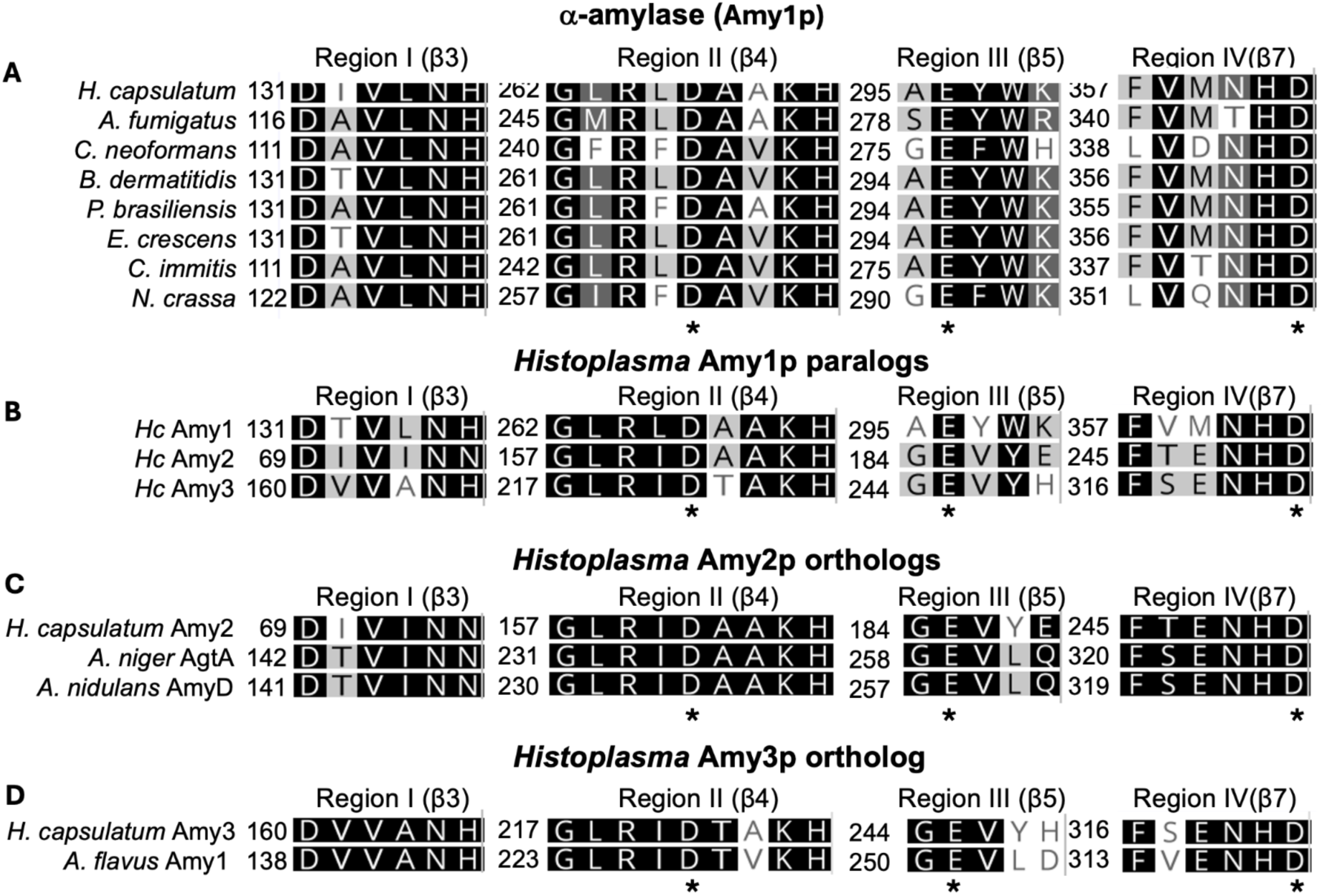
Partial alignment of α-amylases proteins in medically important human fungal pathogens. Amino acid alignment of the four conserved regions comprising the catalytic site of the α-amylase GH13 family. **A.** Alignment between the *H. capsulatum* Amy1p and its homologues in *A. fumigatus, C. neoformans, B. dermatitidis, P. brasiliensis, E. crescens,* and *C. immitis,* and the non-pathogen *N. crassa*. **B.** Alignment of the three paralogs of Amy1p in *Histoplasma capsulatum* (*Hc*). **C.** Alignment of the two orthologs of *Histoplasma capsulatum* Amy2p in *A. niger* AgtA, and *A. nidulans* AmyD. **D.** Alignment of the ortholog of *H. capsulatum* Amy3p in*, A. flavus* Amy1. Conserved α-amylase catalytic residues are shown with an asterisk in all panels.

None of the three Amys in *Histoplasma* showed evidence of having signal peptides, indicating the three proteins are unlikely to be excreted. Amy2p is predicted to have a transmembrane domain at L473 and 2 GPI-anchor signals at S441 and G442, but is predicted to have no secretion signal (Figure S2A-C). Since there is no signal peptide secreted, Amy2p is likely to be synthesized in the cytosol and could be anchored to the membrane. Amy2p’s ortholog in *A. nidulans,* AmyD, has a GPI-anchor site, and it is a membrane-associated protein (He et al. 2017). Overall, these results indicate that even though the catalytic residues are conserved on the three proteins, the cellular location of Amy2p differs from the other two copies.

Next, we searched for *AMY* copies in 19 fungal genomes with the goals of identifying orthologs for *AMY2* and *AMY3*, and establishing how many *AMY* copies existed across different fungi. We found fungi with no copies of *AMY,* like *Saccharomyces cerevisiae,* which was expected, as it does not produce α-glucan. All Amy1 α-amylases belong to the GH13_5 subfamily, while orthologs related to *Histoplasma*’s Amy2p and Amy3p belong to the GH13_1 subfamily. Consistent with previous reports, we found no copies of GH13_32 amylases in ascomycetes, which seem to be restricted to mushroom-body-forming basidiomycetes ((Da Lage et al. 2013), Figure 4A, Figure S1). Among the dimorphic fungi we evaluated, *P. brasiliensis* and *C. immitis* both have one copy of *AMY1* and one copy of *AMY2,* but no copy of *AMY3,* while *B. dermatitidis* has one copy of each, just like *H. capsulatum.* We also found fungi with more than three copies of *AMY,* with *A. niger* having the most out of all the species we analyzed (eight copies). This is consistent with previous evidence showing that the number and type of α-amylases vary depending on the fungal species (reviewed in (Yoshimi et al. 2017)). Phylogenetic analyses of the three paralogs revealed that the *AMY* duplications are ancient (Figure 4B). As expected, *AMY1* is present in both ascomycetes and basidiomycetes. *AMY2* and *AMY3* are also present in the two taxonomic classes. *Histoplasma*’s Amy2p is an ortholog of *A. niger’s* AgtA (an ortholog of *A. nidulans* AmyD), which has been described to be a glucanotransferase that acts on α-1,4-glycosidics bonds (Van Der Kaaij, Yuan, et al. 2007).The full-length protein sequences of these three orthologs are conserved: Amy2p and AgtA show a 54.17 % similarity, Amy2p and AmyD show a 50.21 % similarity, and AgtA and AmyD show a 70.08 % similarity. Alignment and comparison of the four conserved regions (Figure 2C), showed the identity between Amy2p and AgtA as well as Amy2p and AmyD increased to 84.62 %, while the percent identity increased to 100 % in the case between AgtA and AmyD. When we examined a subsample of nine Amy2p orthologs, we found differences in the cellular location of Amy2p orthologs across fungi. None of the assayed orthologs have signal peptides, three of them had six residue omega-sites–including *H. capsulatum* Amy2p–, and two had D residue omega-sites (Table S4). Only four orthologs, *H. capsulatum* Amy2p and three copies in the *A. nidulans*’ genome (AmyC, AmyD, and AmyE, Table S4) had a predicted transmembrane domain (Table S4). The latter is consistent with microscopy observations (He et al. 2017). A similar analysis for the amy3p orthologs revealed that the Amy3p ortholog in *Histoplasma* is the only allele that lacks a signal peptide, a secondary loss that took place since *Histoplasma* diverged from its most common recent ancestor, *Blastomyces* (Table S5). One of the *Histoplasma* Amy3p orthologs is Amy1 from *A. flavus*, which has been implicated in aflatoxin production (Fakhoury and Woloshuk 1999; Gilbert et al. 2018). Comparison of the full length-protein sequences shows 59.48 % similarity, and alignment and comparison of the four conserved regions (Figure 2D), showed the identity increased to 84.62%.

**FIGURE 3.**
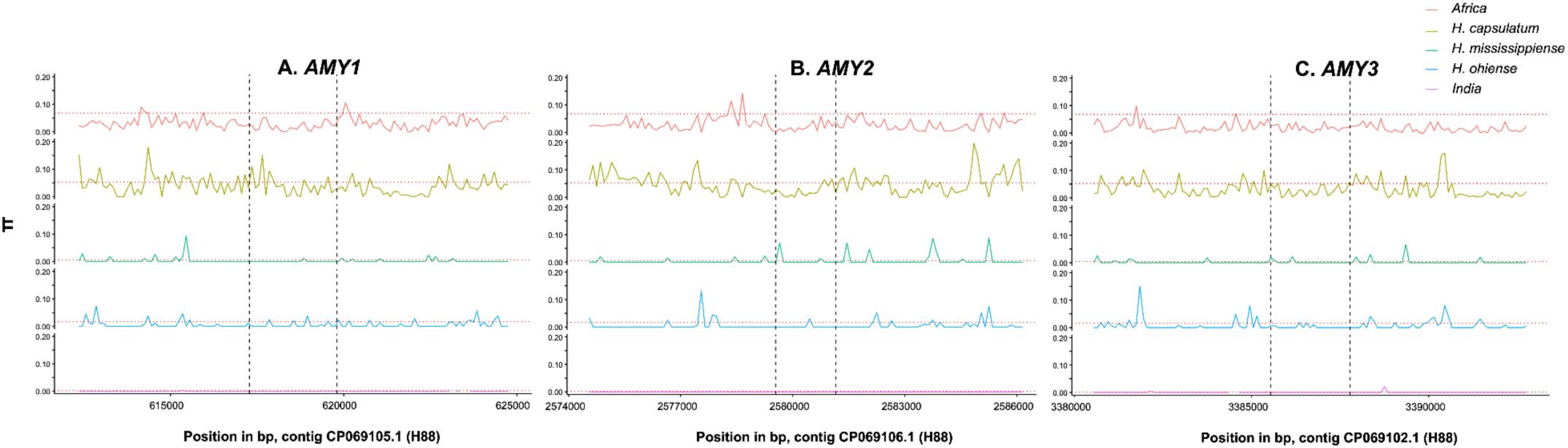
Nucleotide variation (π) in the three *AMY* copies in five phylogenetic species of *Histoplasma*. The red dashed lines show the genomewide average for each species. **A.** *AMY1*. **B.** *AMY2*. **C.** *AMY3*. In general, the three genes show a π lower than the genome average.

**FIGURE 4.**
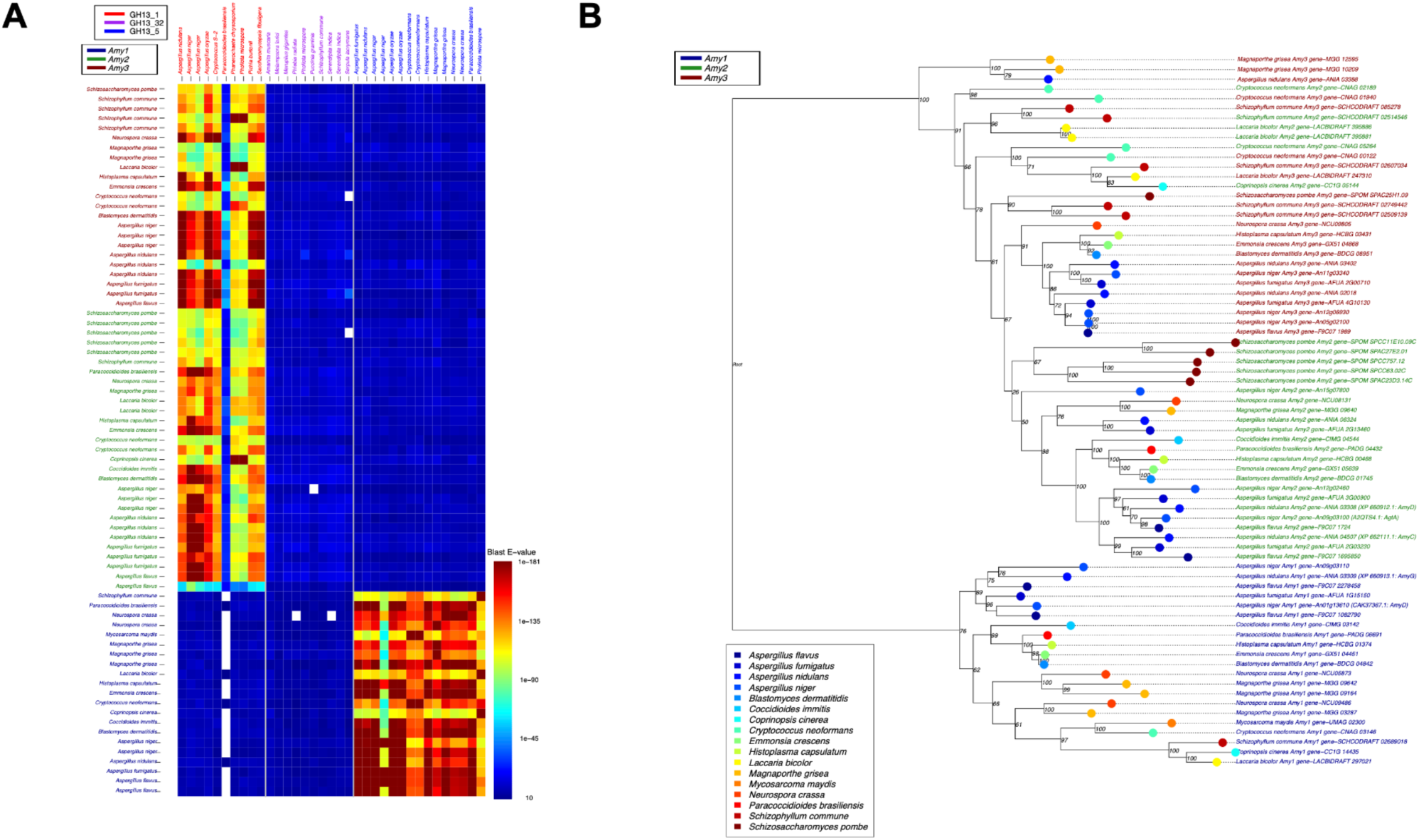
α-amylases found in 19 species of fungi. **A.** Similarity, measured as E-values, of the α-amylases found in Ascomycetes and Basidiomycetes. The color on the heatmap shows the extent of similarity between alleles. The three *H. capsulatum AMY* genes products belong to two different subfamilies of α-amylasesglycoside hydrolases. The color of each cell represents the BLAST e-value when compared with a representative sequence of the three known GH subfamilies. **B.** Maximum likelihood tree of the *AMY* genes in Ascomycetes and Basidomycetes. The labeling (Amy1, Amy2, and Amy3) corresponds to the similarity to the *H. capsulatum ss* genes. Numbers on each node are the bootstrap values.

We ruled out that *Histoplasma*’ *AMY2* and *AMY3* are recent duplicates, as the genes do not appear closely related in the phylogeny. On the contrary, their divergence time appears to predate the split between *Histoplasma* and *Aspergillus*. Note that all *AMY1* orthologs form a well-defined monophyletic clade, whereas the distinction between *AMY2* and *AMY3* orthologs is less clear. The phylogenetic tree shows two monophyletic clades, one corresponding to *AMY2* and one to *AMY3*, but the relationships are not as well resolved as within *AMY1*. For example, the *AMY3* clade includes two genes that are more similar to *AMY2* (*C. neoformans* and *Coprinopsis cinerea*). Moreover, the tree revealed three clades in which *AMY2* and *AMY3* orthologs appear intermingled. The majority of sequences in these three lineages represent basidiomycetous species, but they also include genes from *Magnaporthe grisea* and *A. nidulans*, two ascomycetes.

### Colony morphology

We generated gene disruptions for the three *AMY* copies and *AGS1*. Regardless of the type of guide RNA used to generate mutants, all *ags1-Δ* and *amy1-Δ* strains show a smooth colony morphology, and are consistent with the results previously published of *AGS1* and *AMY1* mutants generated using different methods (Figure 5B) (Rappleye et al. 2004; Marion et al. 2006). Our image analysis quantification of colony roughness also supports the qualitative assessment of colony morphology, but provides more nuanced results (Figure 6). The unscaled mean entropy values ranged from 4.32 - 4.53 among the four analyzed images; after z-score standardization the mean entropy of wild-types across all images was 0 ± 0.92. We found a significant difference between the roughness (measured as pixel entropy) of wild-type colonies and that of the *ags1-Δ* (scaled mean = -1.71 ± 1.83; Wilcoxon rank sum test, *P* < 0.0001) or *amy1-Δ* (scaled mean = -0.80 ± 1.05; Wilcoxon rank sum test, *P* < 0.0001) mutants. The elimination of *AMY2* resulted in a smoother (but not completely smooth) colony morphology, and this was also supported by image analysis. The scaled pixel entropy value for the *amy2-Δ* colonies was -0.25 ± 0.97, greater than what we found for either *ags1-Δ* or *amy1-Δ*, but less than the entropy of the wild-type (Wilcoxon rank sum test, all *P* < 0.0001). Unlike the *amy1-Δ* and *ags1-Δ* strains, the abrogation of *AMY3* did not alter colony morphology, suggesting that *AMY3* is not required for α-(1,3)-glucan production while *AMY2* is partially required. Pixel entropy showed a significant difference between colony roughness of the *amy3-Δ* mutant and the wild-type, but in this case the mutant had greater entropy - and therefore greater roughness - than the wild-type (scaled mean = 0.19 ± 1.03; Wilcoxon rank sum test, *P* < 0.0001). When we standardized the images of *ags1-Δ* and the three *AMY* mutants, we found a gradient of colony roughness with *ags1-Δ* being the least rough, and *amy1-Δ* being slightly greater. *amy3-Δ* produces the roughest colonies, slightly greater than the wild type. *amy2-Δ* produces an intermediate phenotype. There were no colony differences between mutants induced with sgRNA or with tgRNA (Figure 5B).

**FIGURE 5.**
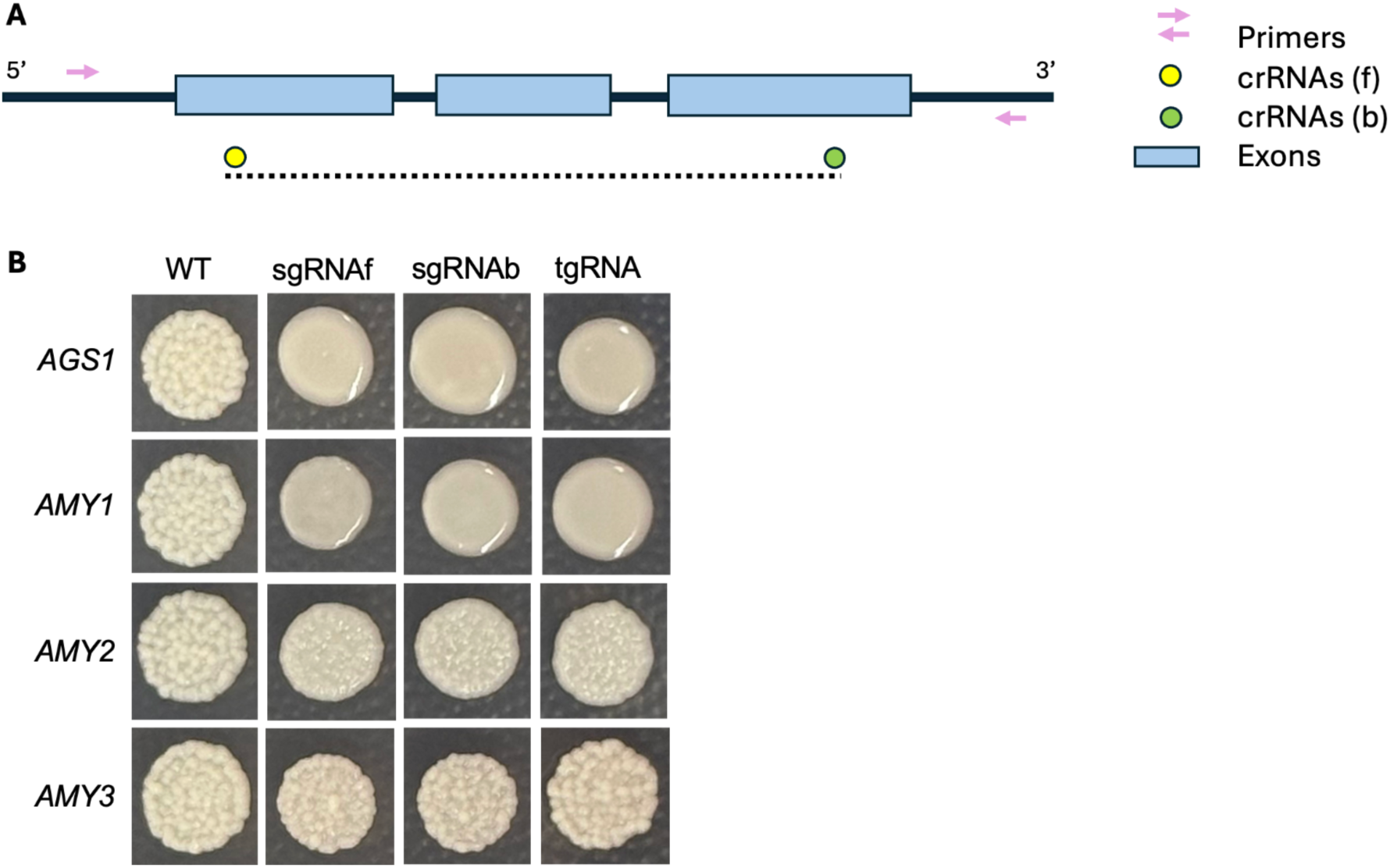
Role of *AGS1*, *AMY1*, *AMY2,* and *AMY3* in the production of α-(1, 3)-glucan. **A.** Schematic representation of the protospacers sequences of the CRISPR RNA (crRNAs, represented by circles) location in the targeted gene (f, front (towards 5’CDS); b, back (towards 3’ CDS)). The dotted line indicates the genomic region deleted using tandem guide RNAs (tgRNA). **B.** Colony morphology of WT *Histoplasma* and *ags1-Δ, amy1-Δ, amy2-Δ,* and *amy3-Δ* mutants generated using single guide RNAs (sgRNA) and tg RNA. f, front; b, back.

**FIGURE 6.**
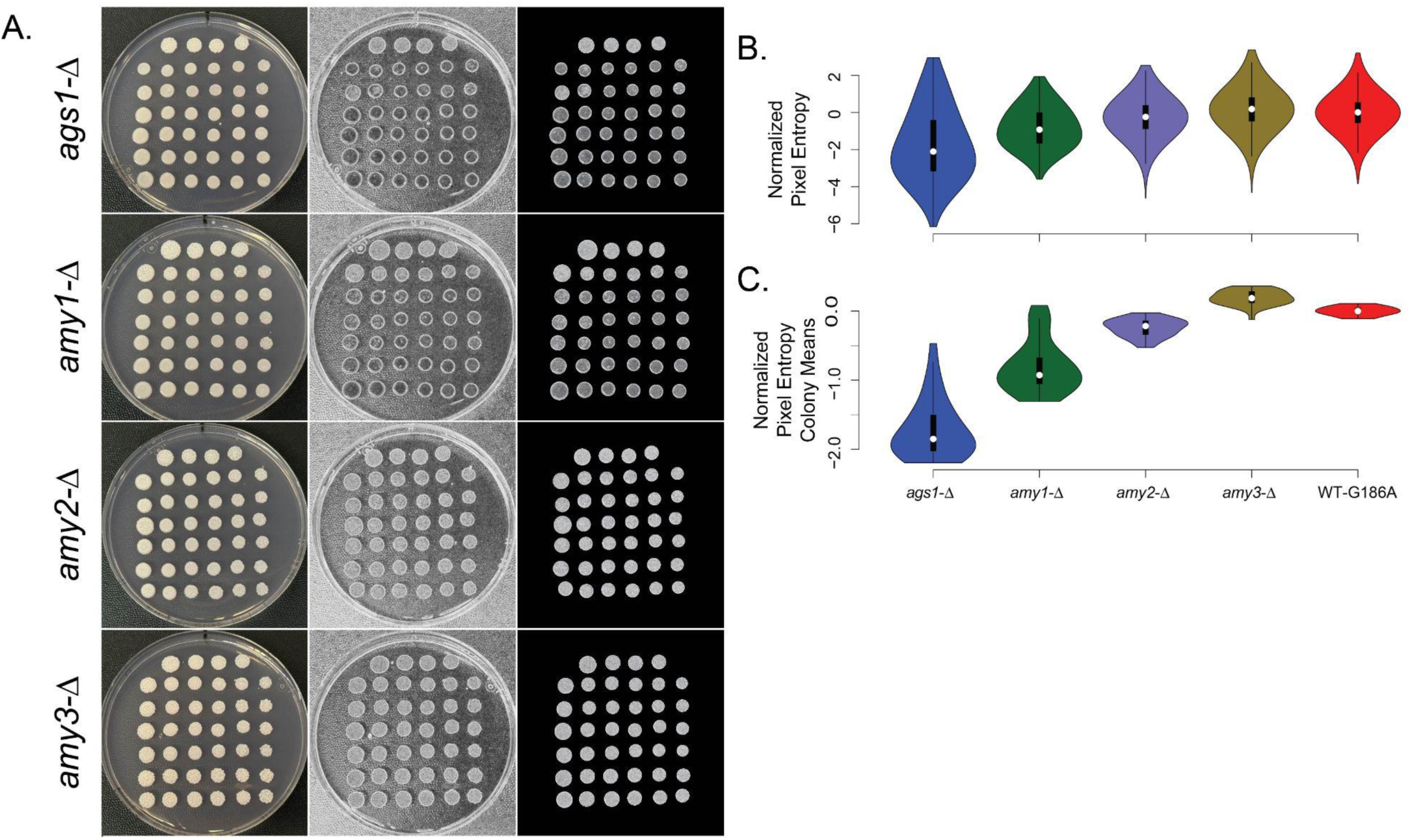
Image analysis quantification of *Histoplasma* colony roughness. **A.** Image analysis workflow showing colony morphology (left column) for 4 mutant strains vs. wild-type. The top four colonies on each plate are wild-type, the remainder are mutants. The middle column depicts the entropy filter applied to grayscale transformations of the colony images. Darker shades show less pixel randomness corresponding with smooth colony morphology; lighter shades show greater entropy and more roughness. The right column shows the result of image segmentation isolating each colony from the image background. **B.** We sampled 1000 random pixels from each segmented colony in the entropy filtered images. Violins show the distributions of these sampled points for each genotype. **C.** Violins show the distribution of colony means for each genotype.

### Growth Curves

We evaluated the rate of growth for the five genotypes by fitting four-parameter logistic models to the growth of each genotype (Figure 7A). These growth curves’ four parameters consisted of the intercept, inflection point, rate of change at the inflection point, and terminal growth. All the growth function curves had similar intercepts as expected by the fact that all growth curves were started with similar inoculum (Figure S3). On the other hand, we detected differences in all the other parameters. All of the points of inflexion, rates of change, and terminal growth differed between the wild type and the mutants (Figure 7B-I). The inflection point differed between all the growth curves (W ≥ 2734, Bonferroni corrected *P* < 0.0001), with the exception of *amy2-Δ* and *amy3-Δ* which showed similar values (W = 5599, Bonferroni corrected *P* = 0.7). Similarly, the rate of change differed among all pairs of genotypes (W ≥ 3, Bonferroni corrected *P* < 0.0001) with the exception of *amy2-Δ* and *amy3-Δ* (W = 5288, Bonferroni corrected *P* = 0.4824). Finally, terminal growth could be described as having three different groups in descending order of growth: WT, *amy2-Δ* and *amy3-Δ*, and *amy1-Δ* and a*gs1-Δ* with the lowest amount of growth after 200 hours. These results indicate that none of the mutants show a growth curve that is similar to the wild-type, but that not all mutants are equal. The results indicate that while *amy2-Δ* and *amy3-Δ have* reduced fitness *in vitro*, they do not show a fitness loss as dramatic as observed in *amy1-Δ* and a*gs1-Δ*.

**FIGURE 7.**
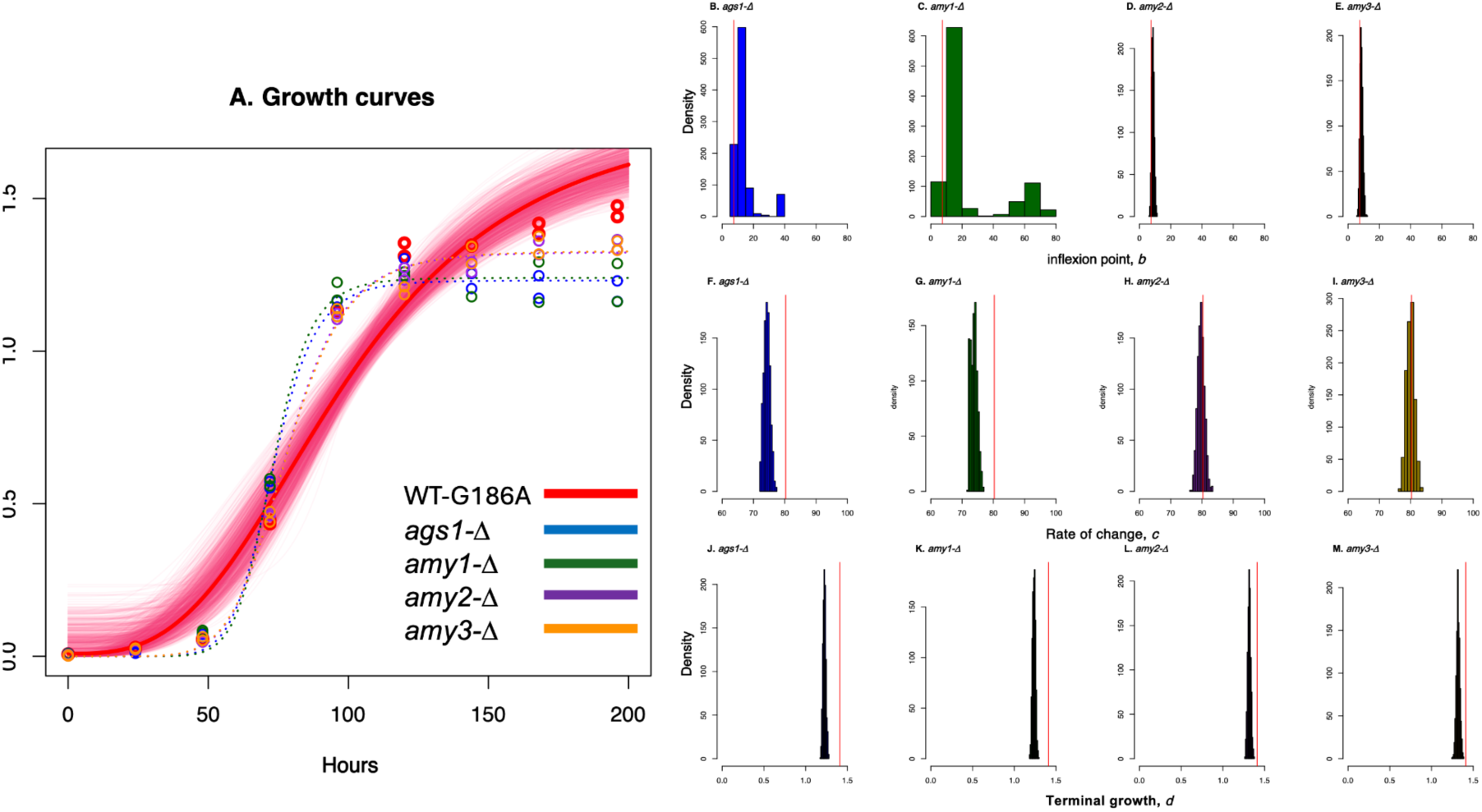
Four-parameter logistic models for the rate of growth of *Histoplasma* (WT and mutants). All experiments were done in HMM broth media at 37°C with 5% CO2. Growth was measured by recording the optical density (OD600) of liquid cultures at different time points (0, 24, 48, 72, 96, 120, 144, 168, 192 hours after inoculation). **A.** Growth curves for the five genotypes. The wildtype, *H. capsulatum ss* G186A (red) is shown with 1,000 regression bootstraps. *ags1-Δ* mutant: blue; *amy1-Δ* mutant: green; *amy2-Δ* mutant: purple; *amy3-Δ* mutant: orange. **B-E**: inflexion point (b) for the fitted curve for each mutant. The red lines show the value of *b* in the wild-type. **F-H**: rate of change (*c*) for the fitted curve for each mutant. The red lines show the value of *b* in the wild-type. **I-L**: terminal growth (*d*) for the fitted curve for each mutant. The red lines show the value of *b* in the wild-type.

### Assessment of the role of *AMY2* and *AMY3* in virulence

Next, we evaluated the mutants’ ability to infect and kill macrophages. We detected strong heterogeneity among mutants generated with tgRNA (F_5,15_ = 20.29, P = 3.428 × 10^-6^) and among mutants generated with sgRNAs (F_5,11_=50.001, P = 3.404 × 10^-7^). Pairwise comparisons among genotypes revealed the source of the heterogeneity. First, we evaluated the virulence of the tg *ags1-Δ* and *amy1-Δ* mutant strains. All *ags1-Δ* and *amy1-Δ* strains were unable to kill the monolayer of macrophages and showed no significant difference in their attenuation when compared to the uninfected control (Figure 8, Table 1). The role of *AGS1* and *AMY1* in *Histoplasma* virulence has been previously described (Rappleye et al. 2004; Marion et al. 2006); these two genes are necessary for the production of α-(1,3)-glucan, and strains in which the genes have been silenced or disrupted fail to kill macrophages. Our experiments using CRISPR modifications confirm these results.

**FIGURE 8.**
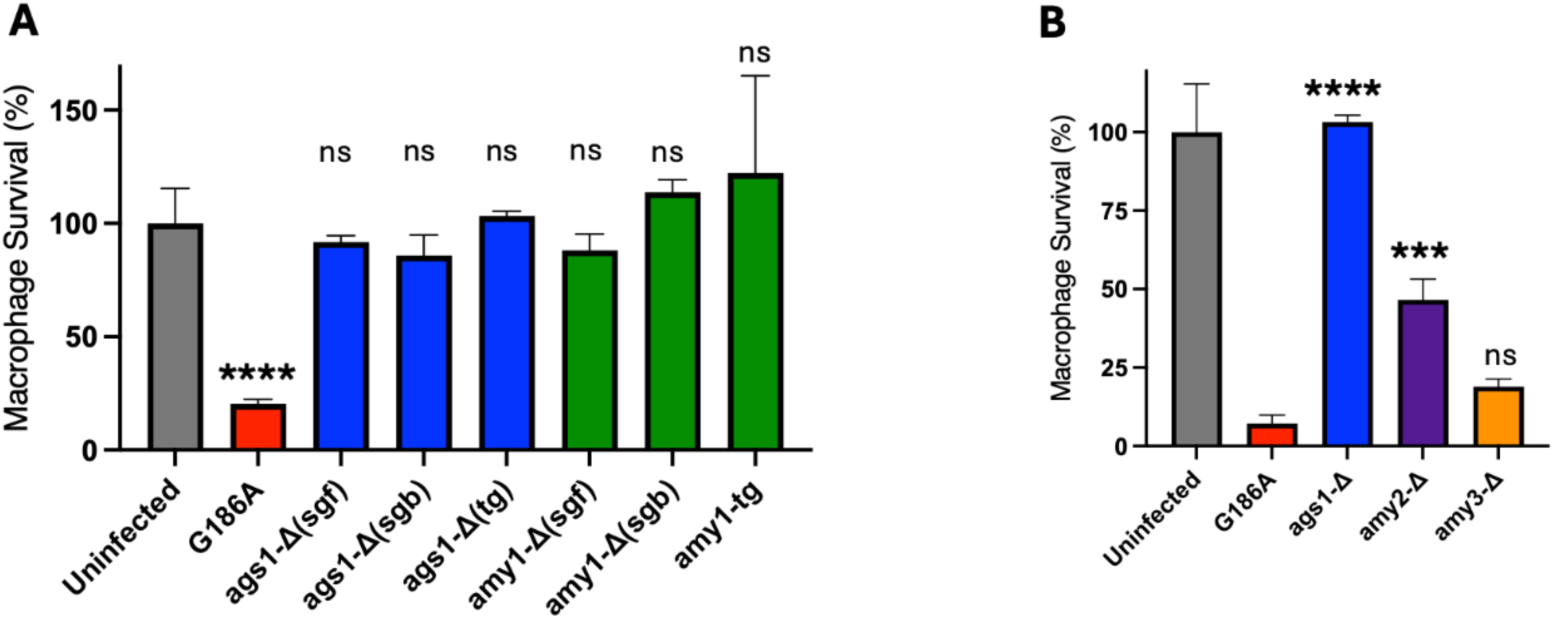
Assessment of *in vitro* virulence of *AMY* mutants. Macrophage survival was measured as the remaining macrophage DNA after coincubation with *Histoplasma* yeast cells. Macrophage DNA remaining at 6 days post-inoculation was normalized to an uninfected population of macrophages. **A.** Virulence comparison of *ags1-Δ* and *amy1-Δ* mutant strains generated using sgRNAs vs tgRNAs. (sgRNA-front (towards 5’CDS): sgf; sgRNA-back (towards 3’CDS), sgb; tgRNA: tg). Data represent results from three independent assays. Error bars represent the standard errors of the means. The one-way ANOVA test was performed to determine statistically significant differences in macrophage survival (%) between each mutant and the uninfected control. ****, P≤0.0001.; ns, not significant. **B.** Role of *AMY2* and *AMY3* in *in vitro* virulence. Lack of *AMY2* partially impairs the virulence of *H. capsulatum* G186A yeast cells in macrophages. Virulence comparison of *ags1-Δ, amy2-Δ* and *amy3-Δ* mutant strains generated by tgRNAs. Data represent results from three independent assays. Error bars represent the standard errors of the means. The one-way ANOVA test was performed to determine statistically significant differences between each mutant and WT. ***, *P* ≤ 0.001; ****, *P* ≤ 0.0001.; ns, not significant.

**TABLE 1.** Tukey HSD pairwise comparisons of virulence levels of mutants generated by tandem guide RNA. THe metric of virulence is remaining macrophages after experimental infections.

| Genotype 1 | Mean<br>genotype 1 | Genotype 2 | Mean<br>genotype 2 | Estimate | Std. Error | t value | P-value |
| --- | --- | --- | --- | --- | --- | --- | --- |
| *amy1*-Δ | 8305.667 | *ags1*-Δ | 7120.667 | 1185.0 | 941.5 | 1.259 | 0.800 |
| *amy2*-Δ | 4484.667 | *ags1*-Δ | 7120.667 | -2636.0 | 815.4 | -3.233 | 0.051 |
| *amy3*-Δ | 1604.667 | *ags1*-Δ | 7120.667 | -5516.0 | 941.5 | -5.859 | < 0.001 |
| G186A | 1483.333 | *ags1*-Δ | 7120.667 | -5637.3 | 941.5 | -5.988 | < 0.001 |
| Uninfected | 7263.000 | *ags1*-Δ | 7120.667 | 142.3 | 941.5 | 0.151 | 0.999 |
| *amy2*-Δ | 4484.667 | *amy1*-Δ | 8305.667 | -3821.0 | 815.4 | -4.686 | 0.003 |
| *amy3*-Δ | 1604.667 | *amy1*-Δ | 8305.667 | -6701.0 | 941.5 | -7.117 | < 0.001 |
| G186A | 1483.333 | *amy1*-Δ | 8305.667 | -6822.3 | 941.5 | -7.246 | < 0.001 |
| Uninfected | 7263.000 | *amy1*-Δ | 8305.667 | -1042.7 | 941.5 | -1.107 | 0.870 |
| *amy3*-Δ | 1604.667 | *amy2*-Δ | 4484.667 | -2880.0 | 815.4 | -3.532 | 0.029 |
| G186A | 1483.333 | *amy2*-Δ | 4484.667 | -3001.3 | 815.4 | -3.681 | 0.022 |
| Uninfected | 7263.000 | *amy2*-Δ | 4484.667 | 2778.3 | 815.4 | 3.407 | 0.037 |
| G186A | 1483.333 | *amy3*-Δ | 1604.667 | -121.3 | 941.5 | -0.129 | 0.999 |
| Uninfected | 7263.000 | *amy3*-Δ | 1604.667 | 5658.3 | 941.5 | 6.010 | < 0.001 |
| Uninfected | 7263.000 | G186A | 1483.333 | 5779.7 | 941.5 | 6.139 | < 0.001 |

We further compared the virulence induced by sgRNA and tgRNA mutations in *AGS1* and *AMY1* to test if the type of guide RNA represented a difference in the ability to kill macrophages. We found no effect of the type of mutation (sg vs tg F_2,12_=2.3194, *P* = 0.1407) or of the interaction of gene under study and the type of mutation (F_2,12_=1.1790, P = 0.3408). These results indicate that mutations introduced by sgRNAs result in the inactivation of a gene just as efficiently as deletions generated by tgRNAs.

Next, we evaluated the *in vitro* virulence of *amy2-Δ* and *amy3-Δ* to test if these two putative α-amylases were also required for virulence in *Histoplasma. amy2-Δ* strain showed a partial attenuation in the ability to kill macrophages. Macrophage viability when infected with *amy2-Δ* was significantly lower than that of uninfected macrophages, or macrophages infected with *amy1-Δ* or *ags1-Δ* but also significantly higher than infections with the wild-type (Figure 8B, Table 1). This result indicates that *AMY2* is necessary for virulence, but its effect size is less pronounced than that of other genes known to affect the production of α-(1,3)-glucan. This also reflects the intermediate (smoother) colony phenotype we observed.

*amy3-Δ* showed no detectable change in its virulence (Figure 8B). Macrophage viability when infected with *amy3-Δ* was similar to the viability when infected with the WT strain. As expected, macrophages infected with *amy3Δ* showed significantly lower viability than the other three mutants, which did not affect viability. These results indicate that *AMY3* is not necessary for virulence, which is consistent with the lack of a detectable effect on colony morphology.

## DISCUSSION

The molecular structure of the cell wall is fundamental to the biological processes and ecological interactions of fungi (Gow et al. 2017). The fungal cell wall is largely composed of polysaccharides such as chitin, β-glucan, and galactomannan, which form a rigid outer layer that determines the interactions between pathogenic fungi and host cells. The specific composition of the cell wall varies at the levels of fungal species, growth stages, and morphological forms. *Histoplasma* grows in its saprophytic mold phase at ∼25°C, and when the temperature shifts to 37°C inside a mammalian host, the morphological transition to its parasitic yeast phase takes place. The cell wall composition is one of the main differences between *Histoplasma* phases. The polysaccharide α-(1,3)-glucan is unique to the yeast phase in most species of *Histoplasma*. The mold phase is characterized by higher content of mannose and lower content of chitin and *β*-glucan (Domer et al. 1967; Domer 1971; Kanetsuna et al. 1974; Reiss et al. 1977a; Reiss et al. 1977b). α-(1,3)-glucan is a key determinant of virulence in the species of *Histoplasma* defined as chemotype II strains (i.e., strains that synthesize α-glucan, (Klimpel and Goldman 1987; Klimpel and Goldman 1988; Eissenberg et al. 1997)). In this piece, we identified two new putative α-amylases belonging to the GH13_1 subfamily in *Histoplasma*, which we dubbed *AMY2* and *AMY3*. We also show that *AMY1* is the only α-amylase encoded by *H. capsulatum* that is fully required for cell wall α-(1,3)-α-glucan biosynthesis, while *AMY2* is partially necessary, and *AMY3* is dispensable. Deletion of the *AGS1* and *AMY1* genes resulted in a smooth colony morphology and complete attenuation of virulence, which is consistent with previous studies (Rappleye et al. 2004; Marion et al. 2006). Deletion of the *AMY2* gene resulted in an intermediate phenotype, while deletion of the *AMY3* gene affected neither the presence of α-(1,3)-glucan in the yeast cell wall nor *in vitro* virulence. Our findings have implications for the understanding of virulence in *Histoplasma* compared to other pathogens, the identification of previously unidentified virulence factors, and the role of gene family evolution in virulence. We discuss each of these implications as follows.

Our experiments confirmed the implication of *AGS1* and *AMY1* in colony morphology and virulence in *Histoplasma*, two genes previously known to affect the production of α-(1,3)-glucan (Rappleye et al. 2004; Marion et al. 2006). α-(1,3) synthase (*AGS1*) was the first gene identified to be essential for α-(1,3)-glucan production in *Histoplasma* (Rappleye et al. 2004). *Histoplasma*’s Amy1p was the first described member of the GH13_5 subfamily of α-amylases, but the orthologs have been characterized in a variety of fungi. The *A. niger* ortholog (AmyD), is involved in synthesizing the primers α-(1,4)-linked glucooligosaccharide, which are required to create longer polymers and stabilize α-(1,3)-glucan (He et al. 2017). In *P. brasiliensis*, Amy1p generates short oligosaccharides that act as primers at the very first step of α-(1,3) glucan production, and has higher hydrolyzing activity on amylopeptin than starch (Camacho et al. 2012). The family includes homologs from *Cryptococcus flavus, C. neoformans, M. grisea, Aspergillus oryzae, A. fumigatus, A. nidulans, A. niger*, and *P. brasiliensis* that were shown to be required for production of α-(1,3)-glucan (Iefuji et al. 1996; Van Der Kaaij, Yuan, et al. 2007; Galdino et al. 2008; Fujikawa et al. 2009; Camacho et al. 2012; He et al. 2017; Kazim et al. 2021). All these orthologs have a conserved structure of catalytic domains and residues. While the pathway involved in the production of α-(1,3)-glucan is dynamic, the roles of Ags1p, and Amy1p in maintaining cell wall structural integrity is generalized in fungi and contribute to virulence in a variety of distant fungal pathogens.

Our analyses also indicate that a second glycoside hydrolase, Amy2p, is also involved in virulence. The abrogation of *AMY2* leads to changes in colony morphology and decreases in macrophage survival rate in the strain G186A, the representative strain of *Histoplasma capsulatum sensu stricto* but its effect is not as large as the one observed when *AGS1* and *AMY1* are disrupted. Of note, Amy2p and Amy3p also have the same catalytic residues present in Amy1p, indicating that there has been selection to maintain the functional features of these paralogs despite millions of years since divergence. On the other hand, Amy2p has both an GPI anchor and a Transmembrane domain which makes it different from Amy1p. This latter feature, the TM domain, makes *Histoplasma*’s Amy2p different from its orthologs in its closest relatives (*Paracoccidioides*, *Coccidioides*) as these alleles do not have that type of domain. Of note, there is another fungus with a similar GPI anchor and a Transmembrane domain: *A. nidulans*. Our phylogenetic analyses indicate that *A. nidulans*’ *AmyD* and *H. capsulatum AMY2* alleles are not each other’s closest relative arguing against horizontal gene transfer. The TM domain and the GPI signature must have evolved through parallel independent evolution in these two genes.

The case of Amy3 is intriguing. *Histoplasma*’s Amy3 differs from other orthologs as the only ortholog without a signal peptide. Moreover, while several ascomycetes species harbor orthologs of this gene, two of the closest relatives of *Histoplasma* do not, *P. brasiliensis* and *C. immitis.* Gene duplications can be resolved through processes like nonfunctionalization, where one copy becomes a pseudogene or is lost (Kimura and Ohta 1974; Ohta 1988; Ohta 1994; Evolution by Gene Duplication | SpringerLink), or through subfunctionalization, where the original gene’s function is partitioned between the two copies. A third possibility is neofunctionalization, in which one copy maintains the original function and the other duplicated gene evolves an entirely new function, and diverges in its effect from the parental copy. We cannot rule out subfunctionalization or neofunctionalization from our data. Nonetheless, we can rule out nonfunctionalization. The *AMY3* gene has been highly conserved across millions of years without degeneration; the conservation of the catalytic domains and residues is particularly notable. The *Histoplasma AMY3* ortholog in *A. flavus* (*Amy1*), encodes an extracellular α-amylase required for aflatoxin production and growth in maize kernels (Fakhoury and Woloshuk 1999; Gilbert et al. 2018). From our data, we can conclude only that the gene is not involved in virulence in mammals. Further, because the *Histoplasma* life cycle includes a probiotic stage, we cannot rule out the possibility that this gene provides fitness advantages in a different life history stage. One possibility is that Amy3 is involved in a different pathway or has undergone neofunctionalization.

Phylogenetic studies have revealed that *Histoplasma capsulatum* is composed of multiple genetically distinct species (Kasuga et al. 1999; Kasuga et al. 2003; Sepúlveda et al. 2017; Maxwell et al. 2018; Jofre et al. 2022; Sepúlveda et al. 2024 May 21). While α-glucan is an important virulence factor for a variety of fungi, there is evidence that the evolution of the pathways involving the production of *β*-glucan and the cell wall composition in *Histoplasma* is evolutionary dynamic. First, α-(1,3)-glucan is required for virulence in all species of *Histoplasma,* except *H. ohiense* (Rappleye et al. 2004; Edwards et al. 2011; Sepúlveda et al. 2014). In this latter species, colonies are smooth, and this phenotype has been attributed to the insertion of transposable elements in the promoter of *AGS1*. Yet *AMY1,* which is downstream of *AGS1,* is still expressed in *H. ohiense* (Edwards et al. 2011) indicating that the downstream components of the pathway are still functional despite not being involved in virulence. These results might indicate that the TE insertions in *AGS1* are recent enough that the pathway has not degenerated but it might do so in the future, or that *AMY1* and potentially other components that remained expressed have pleiotropic effects not involved in the production of α-glucan. Second, and even though α-glucan is important for the cell wall across fungi, its precise proportional representation in the cell wall differs across species. Infrared spectroscopy indicated the presence of an alkali-soluble α-polysaccharide in the cell wall of *H. capsulatum* var. *farciminosum*, a *Histoplasma* species that causes epizootic lymphangitis in horses (Al-Ani 1999; Mapengo et al. 2025). Unlike what has been reported for other species of *Histoplasma* (Kanetsuna et al. 1974), *P. brasiliensis* (Kanetsuna and Carbonell 1970), and *A. niger* (Stagg and Feather 1973), this polymer was not an α-(1,3)-glucan. Instead, the compound is a distinct alkali-soluble polymer that accounts for approximately 45% of the yeast-phase cell wall. This fraction consisted of 30% glucose and 70% of an unidentified reducing compound (San-Blas and Carbonell 1974). The chemical structure of this unspecified component is currently unknown. Similarly, cell wall studies in *H. capsulatum* var. *duboisii* reported a “lactic acid” component (15–30%) released after alkaline hydrolysis of cell walls from serologically nonreactive strains of both species (Pine and Boone 1968).

Gene duplication is considered a major generator of biological novelty (Kondrashov et al. 2002; Roth et al. 2007; Magadum et al. 2013; Panchy et al. 2016), and the resolution of the conflict between gene copies after duplication is a pressing question in evolutionary biology (Kondrashov et al. 2002; Kondrashov et al. 2002; Roth et al. 2007; Hahn 2009; Conant et al. 2014). In bacterial and viral pathogens, gene family expansion is often associated with the emergence of new virulence factors (Anderson and Roth 1977; Perdue et al. 1997; Craven and Neidle 2007; Bernabeu et al. 2019; Sanchez-Herrero et al. 2020; Brinkmann et al. 2023; Lefebvre et al. 2024). Gene duplications of virulence factors have also been detected in fungi (Rikkerink et al. 1990; Wapinski et al. 2007; Brunke et al. 2016; Siscar-Lewin et al. 2019; Sasani et al. 2021; Branco et al. 2023; Merényi et al. 2023; Selmecki 2023). The case we see here is not a recent duplication but instead an ancient one. If gene duplication is an important process in the emergence of virulence, then the study of pathogenesis must move from an individual gene perspective to assess the role of gene families in the trait.

## ACKNOWLEDGEMENTS

This work was funded by the National Institute of Allergy and Infectious Diseases of the National Institutes of Health under award number R01AI153523 to DRM. We are grateful to the Matute lab for comments in previous versions of this manuscript.

## SUPPLEMENTARY MATERIAL

**TABLE S1.** Genomes used to identify GH orthologs.

| Species | Source | fasta | gff |
| --- | --- | --- | --- |
| *Aspergillus flavus* | https://www.ncbi.nlm.nih.gov/datasets/genome/GCF_009017415.1/ | GCF_009017415.1_ASM901741v1_genomic.fna | GCF_009017415.1_ASM901741v1_genomic.gff |
| *Aspergillus fumigatus* | https://www.ncbi.nlm.nih.gov/datasets/genome/GCF_000002655.1/ | GCF_000002655.1_ASM265v1_genomic.fna | GCF_000002655.1_ASM265v1_genomic.gff |
| *Aspergillus nidulans* | https://www.ncbi.nlm.nih.gov/datasets/genome/GCF_000011425.1/ | GCF_000011425.1_ASM1142v1_genomic.fna | GCF_000011425.1_ASM1142v1_genomic.gff |
| *Aspergillus niger* | https://www.ncbi.nlm.nih.gov/datasets/genome/GCF_000002855.4/ | GCF_000002855.4_ASM285v2_genomic.fna | GCF_000002855.4_ASM285v2_genomic.gff |
| *Blastomyces dermatitidis* | https://www.ncbi.nlm.nih.gov/datasets/genome/GCF_000003525.1/ | GCF_000003525.1_BD_ER3_V1_genomic.fna | GCF_000003525.1_BD_ER3_V1_genomic.gff |
| *Coccidioides immitis* | https://www.ncbi.nlm.nih.gov/datasets/genome/GCF_000149335.2/ | GCF_000149335.2_ASM14933v2_genomic.fna | GCF_000149335.2_ASM14933v2_genomic.gff |
| *Coprinopsis cinerea* | https://www.ncbi.nlm.nih.gov/datasets/genome/GCF_000182895.1/ | GCF_000182895.1_CC3_genomic.fna | GCF_000182895.1_CC3_genomic.gff |
| *Cryptococcus neoformans* | https://www.ncbi.nlm.nih.gov/datasets/genome/GCF_000149245.1/ | GCF_000149245.1_CNA3_genomic.fna | GCF_000149245.1_CNA3_genomic.gff |
| *Emmonsia crescens* | https://www.ncbi.nlm.nih.gov/datasets/genome/GCA_002572885.1/ | GCA_002572885.1_Emmon_parv_130_genomic.fna | GCA_002572885.1_Emmon_parv_130_genomic.gff |
| *Histoplasma capsulatum* | https://www.ncbi.nlm.nih.gov/datasets/genome/GCF_000150115.1/ | GCF_000150115.1_ASM15011v1_genomic.fna | GCF_000150115.1_ASM15011v1_genomic.gff |
| *Laccaria bicolor* | https://www.ncbi.nlm.nih.gov/datasets/genome/GCF_000143565.1/ | GCF_000143565.1_V1.0_genomic.fna | GCF_000143565.1_V1.0_genomic.gff |
| *Magnaporthe grisea* | https://www.ncbi.nlm.nih.gov/datasets/genome/GCF_000002495.2/ | GCF_000002495.2_MG8_genomic.fna | GCF_000002495.2_MG8_genomic.gff |
| *Mucor circinelloides* | https://www.ncbi.nlm.nih.gov/datasets/genome/GCA_051295215.1/ | GCA_051295215.1_Mcir_P_S15m_1.0_genomic.fna | GCA_051295215.1_Mcir_P_S15m_1.0_genomic.gff |
| *Mycosarcoma maydis* | https://www.ncbi.nlm.nih.gov/datasets/genome/GCF_000328475.2/ | GCF_000328475.2_Umaydis521_2.0_genomic.fna | GCF_000328475.2_Umaydis521_2.0_genomic.gff |
| *Neurospora crassa* | https://www.ncbi.nlm.nih.gov/datasets/genome/GCF_000182925.2/ | GCF_000182925.2_NC12_genomic.fna | GCF_000182925.2_NC12_genomic.gff |
| *Paracoccidioides brasiliensis* | https://www.ncbi.nlm.nih.gov/datasets/genome/GCF_000150735.1/ | GCF_000150735.1_Paracocci_br_Pb18_V2_genomic.fna | GCF_000150735.1_Paracocci_br_Pb18_V2_genomic.gff |
| *Saccharomyces cerevisiae* | https://www.ncbi.nlm.nih.gov/datasets/genome/GCF_000146045.2/ | GCF_000146045.2_R64_genomic.fna | GCF_000146045.2_R64_genomic.gff |
| *Schizophyllum commune* | https://www.ncbi.nlm.nih.gov/datasets/genome/GCF_000143185.2/ | GCF_000143185.2_Schco3_genomic.fna | GCF_000143185.2_Schco3_genomic.gff |
| *Schizosaccharomyces pombe* | https://www.ncbi.nlm.nih.gov/datasets/genome/GCA_000002945.3/ | GCF_000002945.2_ASM294v3_genomic.fna | GCF_000002945.2_ASM294v3_genomic.gff |

**TABLE S2.**
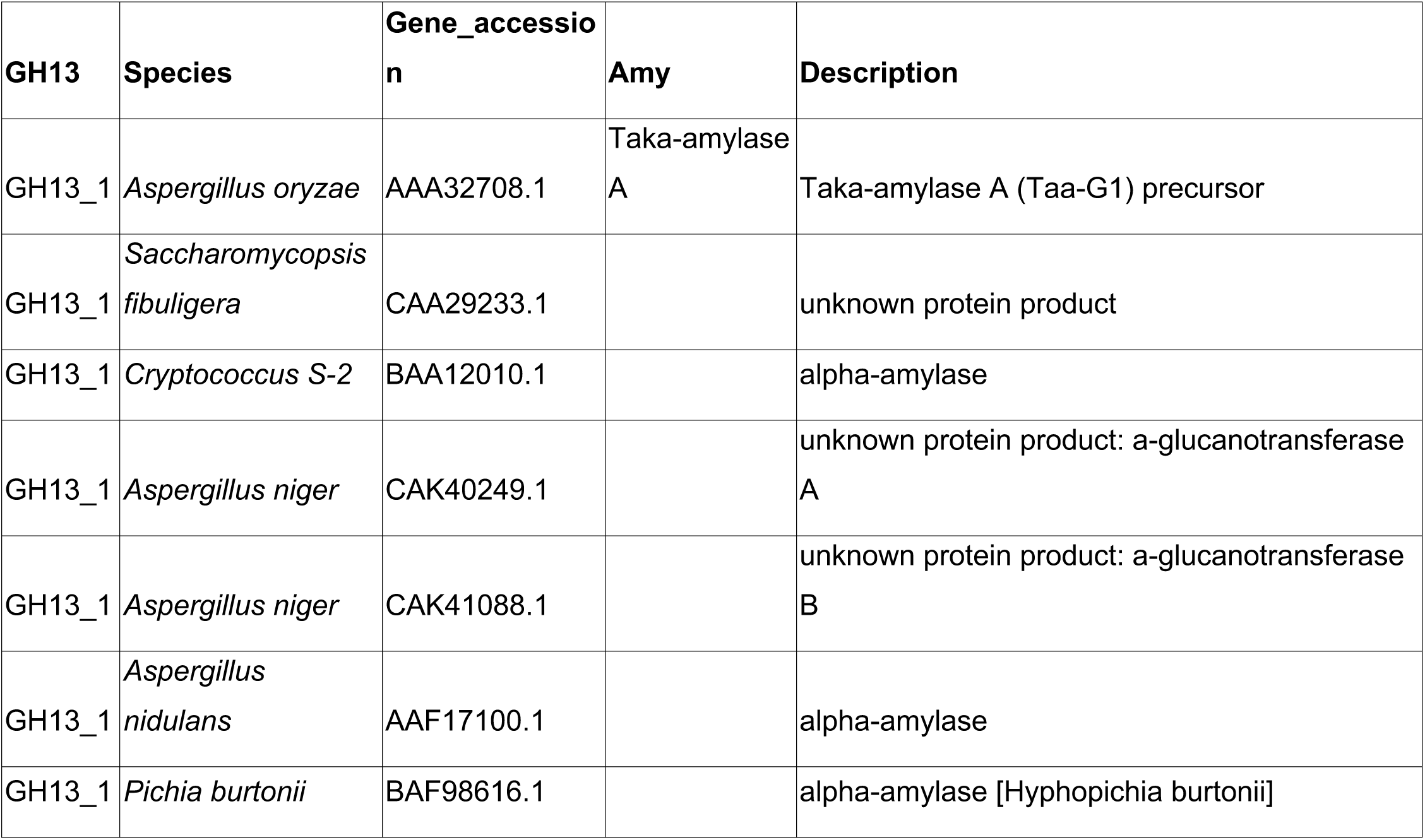
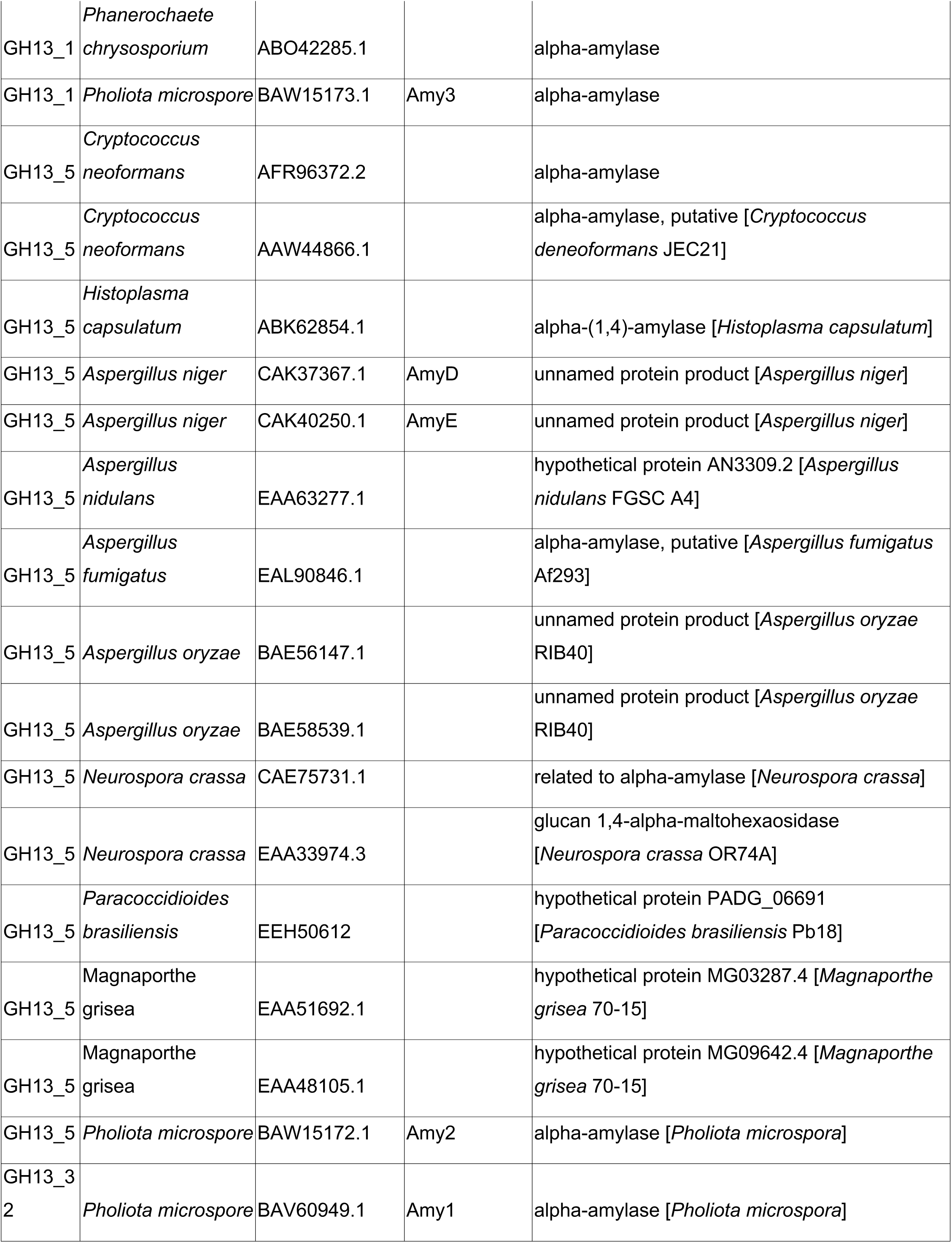
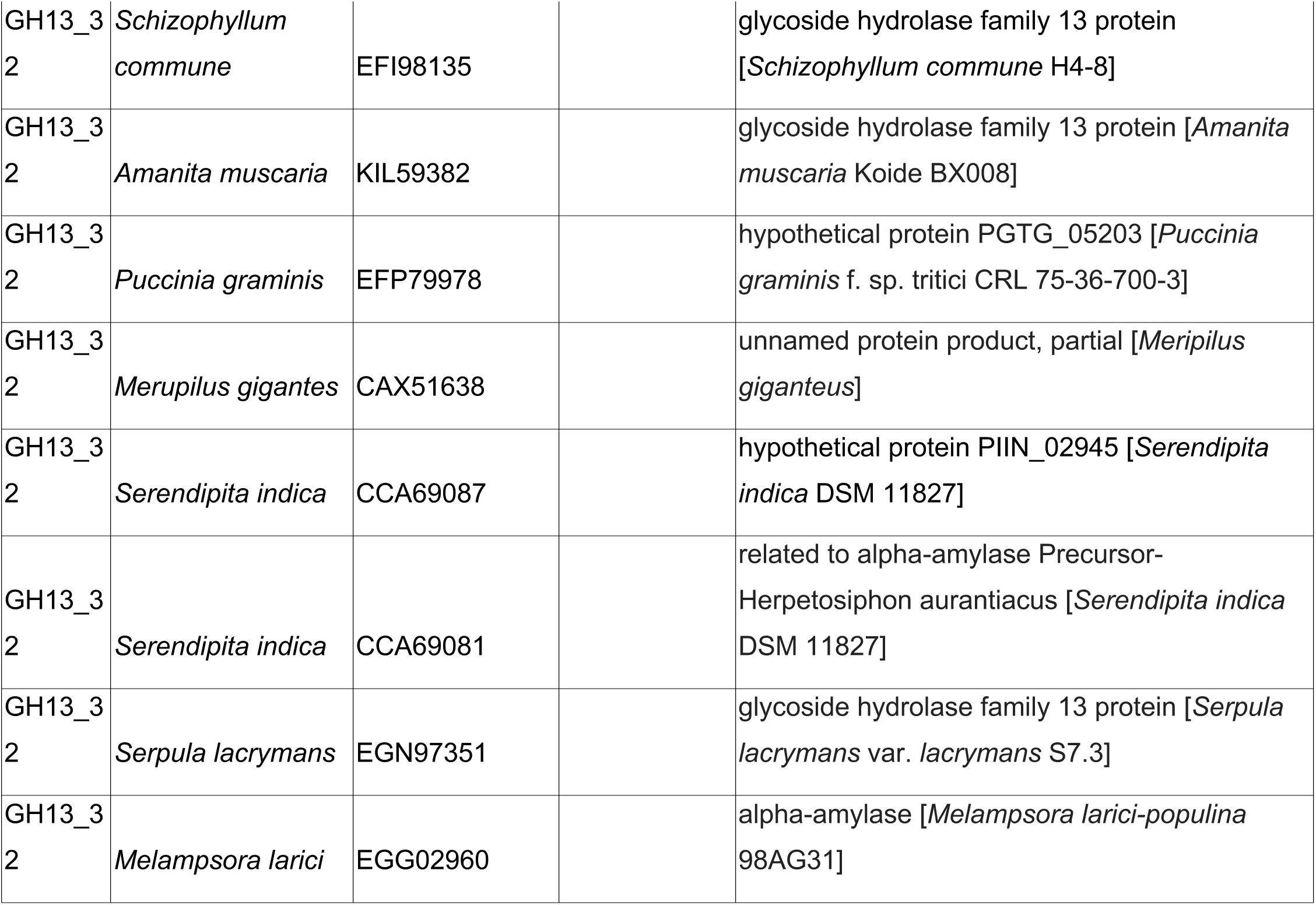
List of known α-amylases belonging to the GH13_1, GH13_5, and GH13_32 subfamilies.

**TABLE S3.**
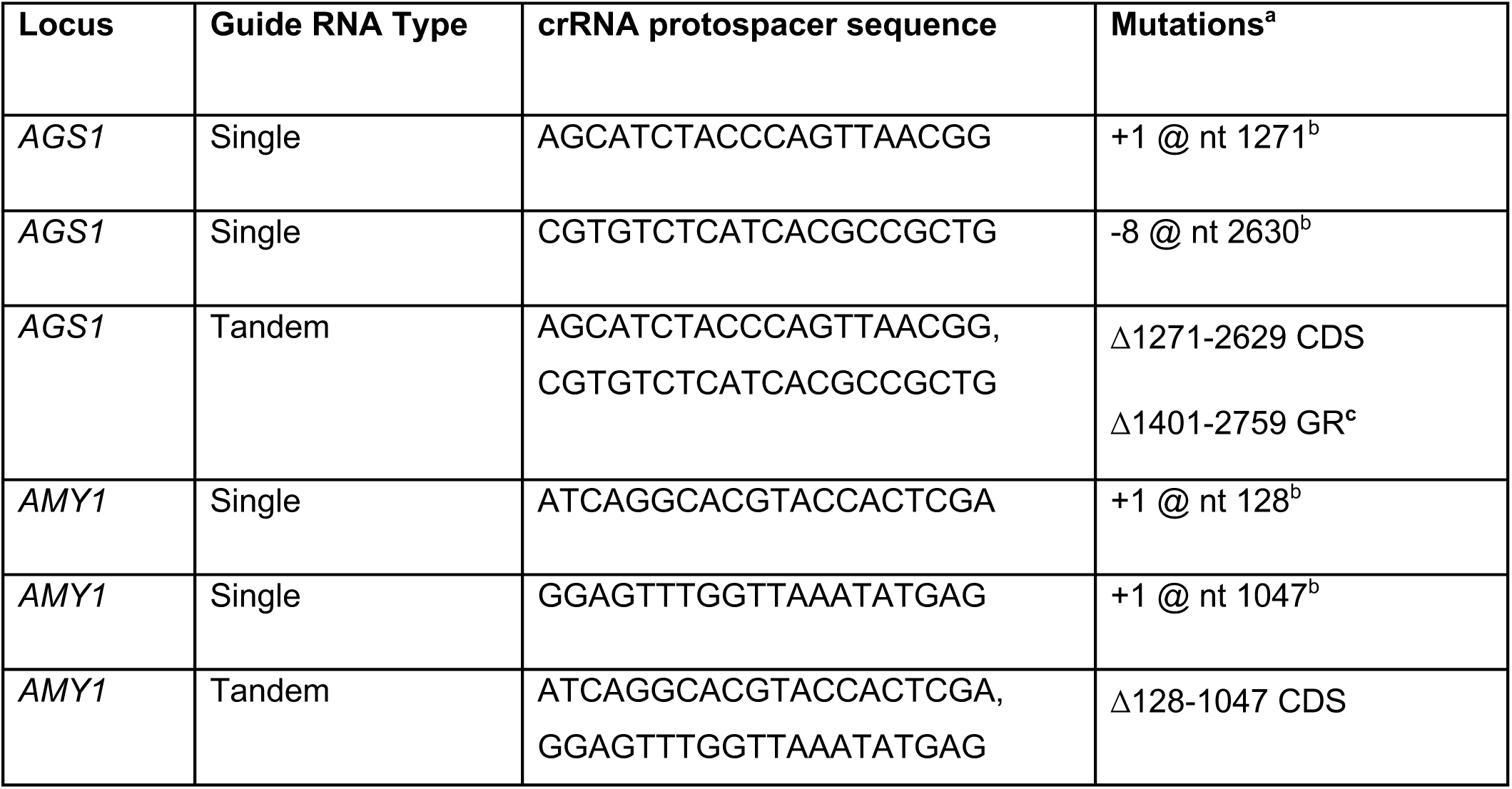
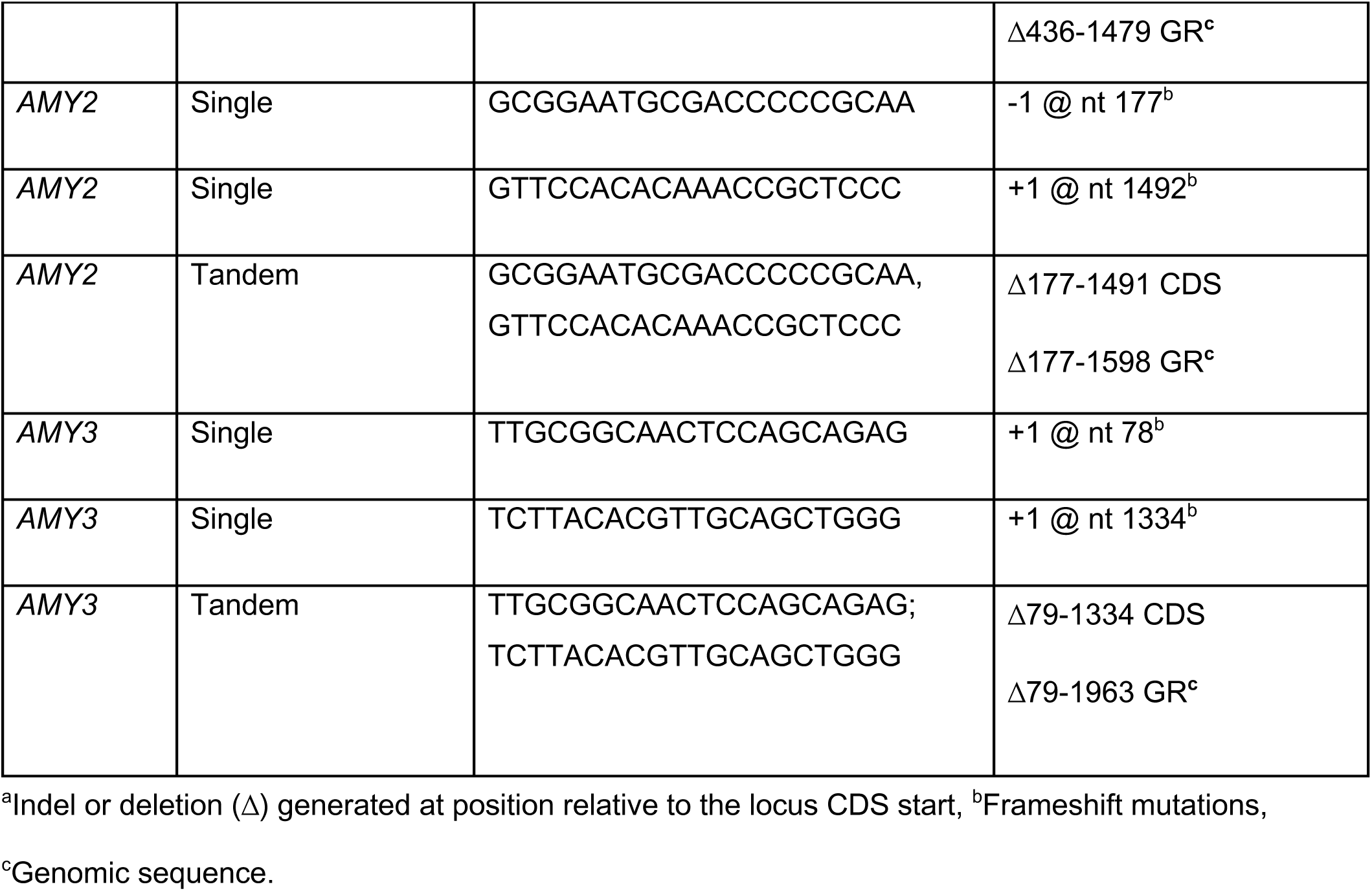
CRISPR/Cas9-generated *Histoplasma* mutant strains.

**TABLE S4.** Signal peptide and transmembrane domain detection in Amy2p orthologs.

| Species | Accession Number (protein) | Gene / Protein Name | Signal peptide? | GPI anchor? | Transmembrane domain? |
| --- | --- | --- | --- | --- | --- |
| *Aspergillus nidulans*<br>FGSC A4 | XP_662111.1 | amyC<br>(amylase C; α-amylase-like) | not found | S residue<br>omega-site<br>predicted at<br>position 491. | 1 TMhelix<br>between 494<br>& 516. |
| *Aspergillus fumigatus*<br>Z5 | KMK57693.1 | AmyA (α-amylase) | Signal peptide:<br>Cleavage site<br>between pos.<br>26 and 27. | S residue<br>omega-site<br>predicted at<br>position 543. | 0 |
| *Aspergillus niger* | A2QTS4<br>.1 | agtA (4-α-glucanotransferase) | Signal peptide:<br>Cleavage site between pos. 25 and 26. | D residue<br>omega-site predicted at position 533. | 0 |
| *Coccidioides immitis*<br>RS | EAS335<br>20.3 | AmyA (α-amylase) | Signal peptide:<br>Cleavage site between pos. 27 and 28. | S residue<br>omega-site predicted at position 556. | 0 |
| *Paracoccidioides brasiliensis*<br>Pb18 | EEH483<br>48.2 | PADG_04432 (hypothetical amylase-like protein) | Signal peptide:<br>Cleavage site between pos. 16 and 17. | S residue<br>omega-site predicted at position 509. | 0 |
| *Cryptococcus neoformans* | XP_012048813.1 | alpha-amylase AmyA, variant | not found | not anchored. | 0 |
| *Schizosaccharomyces pombe* | NP_001342914.1 | alpha-amylase-like protein | not found | not anchored. | 0 |
| *Aspergillus nidulans*<br>FGSC A4 | XP_660912.1 | amyD | not found | S residue<br>omega-site predicted at position 526. | 0 |
| Amy2p Hc<br>(our sequence) |  |  | not found | S residue<br>omega-site predicted at position 443. | 1 TMhelix between 450 & 472. |
| *Blastomyces dermatitidis*<br>ER-3 | EEQ866<br>25.1 | pullulanase | Signal peptide:<br>Cleavage site between pos. 26 and 27. | S residue<br>omega-site predicted at position 523. | 0 |
| *Aspergillus nidulans*<br>FGSC A4 | XP_663<br>928.1 | amyE | Signal peptide:<br>Cleavage site<br>between pos.<br>20 and 21. | N residue<br>omega-site<br>predicted at<br>position 537. | 0 |
| *Aspergillus niger* CBS<br>513.88 | A2QYT9<br>.1 | agtB (4-alpha-<br>glucanotransfe<br>rase) | Signal peptide:<br>Cleavage site<br>between pos.<br>28 and 29. | S residue<br>omega-site<br>predicted at<br>position 522. | 1 TMhelix<br>between 525<br>& 527. |
| *Aspergillus fumigatus*<br>Af293 | XP_748<br>411.1 | alpha-amylase | Signal peptide:<br>Cleavage site<br>between pos.<br>25 and 26. | S residue<br>omega-site<br>predicted at<br>position 526. | 1 TMhelix<br>between 537<br>& 559. |
| *Aspergillus flavus* | XP_041<br>143812.<br>1 | putative alpha-amylase | Signal peptide:<br>Cleavage site<br>between pos.<br>23 and 24. | S residue<br>omega-site<br>predicted at<br>position 529. | 0 |
| *Aspergillus flavus* | QRD84<br>679.1 | glycoside<br>hydrolase<br>superfamily | Signal peptide:<br>Cleavage site<br>between pos.<br>25 and 26. | not anchored. | 1 TMhelix<br>between 7 &<br>26. |
| *Aspergillus fumigatus*<br>Af293 | XP_755<br>679.1 | alpha-amylase | Signal peptide:<br>Cleavage site<br>between pos.<br>20 and 21. | S residue<br>omega-site<br>predicted at<br>position 535. | 0 |
| *Cryptococcus neoformans* H99 | AFR956<br>31.1 | alpha-amylase | Signal peptide:<br>Cleavage site<br>between pos.<br>21 and 22. | S residue<br>omega-site<br>predicted at<br>position 506. | 0 |
| *Schizophyllum commune*<br>H4-8 | KAI5887<br>882.1 | glycoside<br>hydrolase<br>family 13<br>protein | Signal peptide:<br>Cleavage site<br>between pos.<br>20 and 21. | N residue<br>omega-site<br>predicted at<br>position 503. | 0 |
| *Laccaria*<br>*bicolor*<br>S238N-H82 | XP_001<br>888631.<br>1 | GH13 alpha-amylase<br>precursor | Signal peptide:<br>Cleavage site<br>between pos.<br>21 and 22. | S residue<br>omega-site<br>predicted at<br>position 536. | 0 |
| *Laccaria*<br>*bicolor*<br>S238N-H82 | EDR006<br>21.1 | GH13 alpha-amylase<br>precursor | Signal peptide:<br>Cleavage site<br>between pos.<br>21 and 22. | not anchored. | 0 |
| *Coprinopsi*<br>*s cinerea*<br>okayama7<br>#130 | EAU883<br>78.2 | alpha-amylase | Signal peptide:<br>Cleavage site<br>between pos.<br>19 and 20. | S residue<br>omega-site<br>predicted at<br>position 498. | 0 |
| *Neurospor*<br>*a crassa*<br>OR74A | XP_959<br>674.1 | alpha-amylase | Signal peptide:<br>Cleavage site<br>between pos.<br>22 and 23. | N residue<br>omega-site<br>predicted at<br>position 509. | 0 |
| *Schizosacc*<br>*haromyces*<br>*pombe* | NP_594<br>401.1 | glycosyl<br>hydrolase<br>family 13<br>protein | not found | S residue<br>omega-site<br>predicted at<br>position 551. | 0 |
| *Schizosacc*<br>*haromyces*<br>*pombe* | NP_587<br>687.1 | GPI-anchored<br>glycosyl<br>hydrolase<br>family 13<br>protein | Signal peptide:<br>Cleavage site<br>between pos.<br>22 and 23. | S residue<br>omega-site<br>predicted at<br>position 597. | 0 |
| *Schizosacc*<br>*haromyces*<br>*pombe* | NP_587<br>976.1 | GPI-anchored<br>glycosyl<br>hydrolase<br>family 13<br>protein | Signal peptide:<br>Cleavage site<br>between pos.<br>21 and 22. | S residue<br>omega-site<br>predicted at<br>position 538. | 1 TMhelix<br>between 541<br>& 563. |
| *Schizosaccharomyces pombe* | NP_594551.1 | alpha-amylase<br>2 | Signal peptide:<br>Cleavage site<br>between pos.<br>24 and 25. | not anchored. | 1 TMhelix<br>between 7 &<br>26. |

**TABLE S5.** Signal peptide and transmembrane domain detection in Amy3p orthologs.

| Species | Accession Number (protein) | Gene / Protein Name | Signal peptide? | GPI anchor? | Transmembrane domain? |
| --- | --- | --- | --- | --- | --- |
| *Aspergillus nidulans*<br>FGSC A4 | XP_659622.1 | amyA (α-amylase A) | Signal peptide:<br>Cleavage site<br>between pos.<br>15 and 16. | not anchored. | 0 |
| *Aspergillus nidulans*<br>FGSC A4 | XP_661006.1 | amyB (α-amylase B) | Signal peptide:<br>Cleavage site<br>between pos.<br>19 and 20. | not anchored. | 0 |
| *Aspergillus fumigatus*<br>Af293 | XP_077660772.1 | alpha-amylase | Signal peptide:<br>Cleavage site<br>between pos.<br>19 and 20. | not anchored. | 0 |
| *Aspergillus niger* | XP_001395749.3 | extracellular<br>alpha-amylase<br>(amyA/amyB-type) | Signal peptide:<br>Cleavage site<br>between pos.<br>20 and 21. | not anchored. | 0 |
| *Cryptococcus neoformans*<br>H99 | XP_012046544.1 | alpha-amylase | Signal peptide:<br>Cleavage site<br>between pos.<br>22 and 23. | S residue<br>omega-site<br>predicted at<br>position 529. | 0 |
| *Blastomyces dermatitidis* ER-3 | XP_045282744.1 | alpha-amylase A type-3, variant 2 | Signal peptide:<br>Cleavage site between pos. 20 and 21. | not anchored. | 0 |
| *Schizosaccharomyces pombe* | NP_001343001.1 | alpha-amylase | Signal peptide:<br>Cleavage site between pos. 25 and 26. | not anchored. | 0 |
| *Aspergillus flavus* | XP_041143568.1 | putative alpha-amylase | Signal peptide:<br>Cleavage site between pos. 21 and 22. | not anchored. | 0 |
| Amy3p Hc (our sequence) |  |  | not found. | not anchored. | 0 |
| *Aspergillus fumigatus* Af293 | XP_749208.1 | putative alpha-amylase | Signal peptide:<br>Cleavage site between pos. 23 and 24. | not anchored. | 0 |
| *Aspergillus niger* | XP_001394335.1 | acid alpha-amylase-Aspergillus niger | Signal peptide:<br>Cleavage site between pos. 21 and 22. | not anchored. | 0 |
| *Aspergillus niger* | XP_001390778.3 | extracellular alpha-amylase amyA/amyB-Aspergillus niger | Signal peptide:<br>Cleavage site between pos. 20 and 21. | not anchored. | 0 |
| *Cryptococcus neoformans* H99 | XP_012052859.1 | alpha-amylase | Signal peptide:<br>Cleavage site between pos. 25 and 26. | not anchored. | 0 |
| *Aspergillus nidulans*<br>FGSC A4 | XP_660<br>992.1 | amyF | Signal peptide:<br>Cleavage site<br>between pos.<br>20 and 21. | not<br>anchored. | 0 |
| *Schizophyllum commune*<br>H4-8 | KAI5891<br>966.1 | glycoside<br>hydrolase<br>family 13<br>protein | Signal peptide:<br>Cleavage site<br>between pos.<br>20 and 21. | G residue<br>omega-site<br>predicted at<br>position 494. | 0 |
| *Schizophyllum commune*<br>H4-8 | KAI5900<br>115.1 | glycoside<br>hydrolase<br>family 13<br>protein | Signal peptide:<br>Cleavage site<br>between pos.<br>25 and 26. | N residue<br>omega-site<br>predicted at<br>position 510. | 0 |
| *Schizophyllum commune*<br>H4-8 | KAI5891<br>419.1 | glycoside<br>hydrolase<br>family 13<br>protein | Signal peptide:<br>Cleavage site<br>between pos.<br>18 and 19. | not<br>anchored. | 0 |
| *Schizophyllum commune*<br>H4-8 | KAI5890<br>653.1 | glycoside<br>hydrolase<br>family 13<br>protein | Signal peptide:<br>Cleavage site<br>between pos.<br>18 and 19. | not<br>anchored. | 0 |
| *Laccaria bicolor*<br>S238N-H82 | EDR113<br>45.1 | glycoside<br>hydrolase<br>family 13<br>protein | Signal peptide:<br>Cleavage site<br>between pos.<br>19 and 20. | S residue<br>omega-site<br>predicted at<br>position 505. | 1 TMhelix<br>between 510<br>& 532. |
| *Neurospora crassa*<br>OR74A | XP_964<br>065.2 | alpha-amylase | Signal peptide:<br>Cleavage site<br>between pos.<br>23 and 24. | not<br>anchored. | 1 TMhelix<br>between 7 &<br>24. |

**FIGURE S1.**
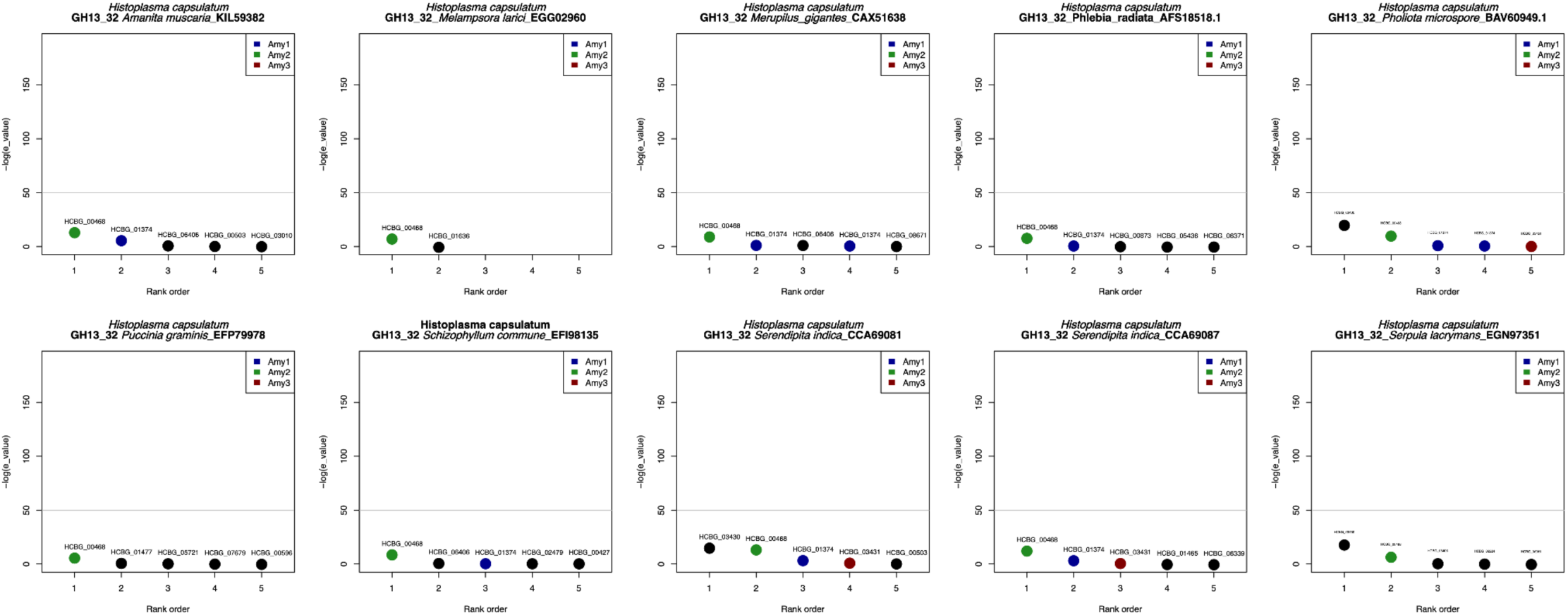
No evidence of GH13_32 amylases in the genome of *H. capsulatum*. Values in the y-axis represent BLAST E-values. The second name in the title refers to the GH13_32 allele compared against the *H. capsulatum* genome.

**FIGURE S2.**
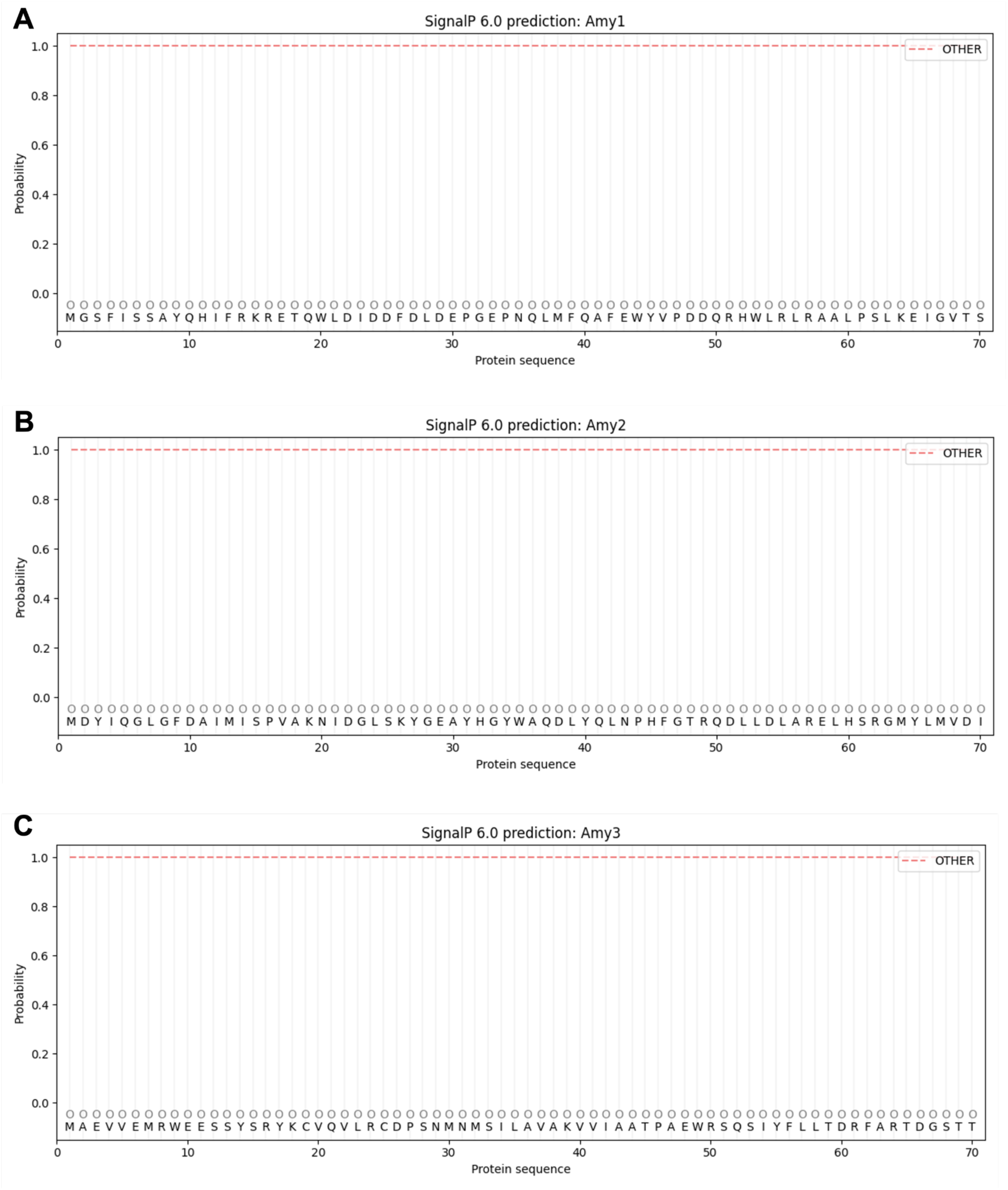
None of the three *Histoplasma* Amy copies has a signal peptide. The plots show the prediction as determined by SignalP 6.0. A. Amy1p. B. Amy2p. C. Amy3p.

**FIGURE S3.**
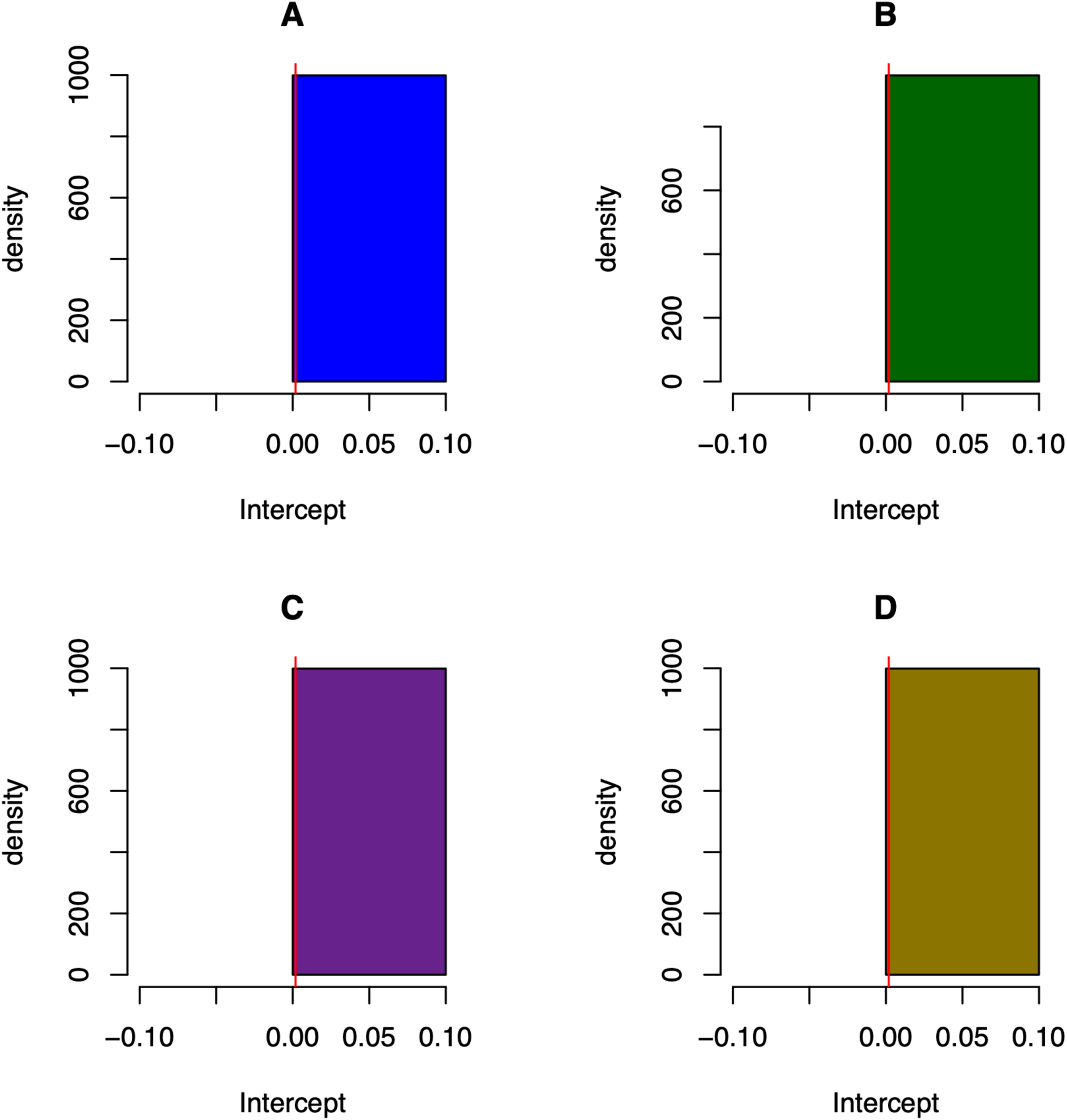
The growth curves of all genotypes show similar intercepts. **A.** *ags1-Δ* mutant. **B.** *amy1-Δ* mutant. **C.** *amy2-Δ* mutant. **D.** *amy3-Δ* mutant. The red line in each panel shows the intercept of the wildtype (*Histoplasma capsulatum sensu stricto* G186A).

